# TXNDC17 and glutathione cooperate to prevent cystine-induced cell death

**DOI:** 10.64898/2026.11.19.767784

**Authors:** Caroline Coelho de Faria, Mami Sato, Shibo Sun, Thamara Nishida Xavier da Silva, Qidong Li, Sebastian Bothe, Gereon Poschmann, Seiryo Ogata, Uladzimir Barayeu, Takaaki Akaike, Gholamreza Fazeli, Elias SJ Arnér, Carsten Berndt, Boyi Gan, José Pedro Friedmann Angeli

**Affiliations:** Rudolf Virchow Zentrum (RVZ), Center for Integrative and Translational Bioimaging, University of Würzburg, Würzburg, Germany; Department of Experimental Radiation Oncology, The University of Texas MD Anderson Cancer Center, Houston, TX, USA; Institute of Molecular Medicine, Proteome Research, University Hospital and Medical Faculty, Heinrich Heine University Düsseldorf, Düsseldorf, Germany; Max-Planck-Institute for Polymer Research, Mainz, Germany; Department of Redox Molecular Medicine, Tohoku University Graduate School of Medicine, Sendai, Japan; Division of Biochemistry, Department of Medical Biochemistry and Biophysics, Karolinska Institutet, Stockholm, Sweden; Department of Selenoprotein Research and the National Tumor Biology Laboratory, National Institute of Oncology, Budapest, Hungary; Department of Neurology, University Hospital and Medical Faculty, Heinrich Heine University Düsseldorf, Düsseldorf, Germany; Comprehensive Cancer Center Mainfranken, University hospital Würzburg, Würzburg, Germany

## Abstract

Glutathione (GSH) supports diverse metabolic and antioxidant functions, yet the cellular processes most vulnerable to its depletion remain incompletely defined. Using genome-wide genetic screening in GSH-depleted cells, we identified the cystine reductase TXNDC17 as essential for survival under GSH limitation. GSH depletion in *TXNDC17*-knockout cells caused intracellular cystine accumulation, increased protein cysteinylation, and cell death. Inhibiting the cystine/glutamate antiporter system x_C_⁻ prevented cell death, linking this vulnerability to cystine uptake and impaired intracellular reduction. GSH depletion also markedly suppressed the growth of xenografts derived from *TXNDC17*-knockout, *KEAP1*-mutant lung cancer cells with high system x_C_⁻ activity, accompanied by reduction-sensitive changes in cytoskeletal proteins. We further identified an electrophilic fragment that induced dose-dependent cytotoxicity in GSH-depleted cancer cells in a cystine-uptake dependent manner. Together, our findings establish TXNDC17 and GSH as central regulators of cystine-driven cytotoxicity, uncovering a therapeutic vulnerability in cancers with high cystine uptake.

---

Glutathione (GSH) is the most abundant low-molecular-weight thiol in mammalian cells and supports multiple aspects of cellular physiology and homeostasis ^1^. It provides reducing equivalents for peroxide detoxification, including reducing phospholipid hydroperoxides via glutathione peroxidase 4 (GPX4), thereby suppressing ferroptosis ^2–5^. GSH also participates in electrophile conjugation, regulates protein function through S-glutathionylation, and contributes to iron metabolism ^6,7^. Beyond these functions, GSH also acts as a mobilizable cysteine reservoir, where intracellular GSH catabolism can sustain mitochondrial respiration during cysteine starvation, whereas extracellular GSH breakdown supplies cysteine to support tumor growth ^8,9^.

Maintaining GSH synthesis requires a sustained cysteine supply. Under standard cell culture conditions, many cells acquire cysteine predominantly through uptake of its oxidized disulfide form, cystine, by the cystine–glutamate antiporter system x_c_⁻. This transporter comprises the substrate-selective light chain xCT, encoded by *SLC7A11*, and the heavy chain 4F2hc, encoded by *SLC3A2* ^10^. System x_c_⁻ couples cystine import to glutamate export, after which intracellular reduction generates cysteine for GSH and protein synthesis (Extended Data Fig. 1a). GSH is synthesized through two ATP-dependent reactions: glutamate–cysteine ligase catalyzes the formation of γ-glutamylcysteine, and glutathione synthetase subsequently adds glycine mitochondrial respiration during cysteine starvation ^11^.

Cystine uptake and GSH synthesis are coordinated in part by the transcription factor NRF2, which promotes expression of *SLC7A11* and the catalytic and modifier subunits of glutamate–cysteine ligase, *GCLC* and *GCLM* ^12^. Under basal conditions, KEAP1 promotes NRF2 ubiquitination and proteasomal degradation, whereas oxidative or electrophilic stress impairs this process, allowing NRF2 to accumulate and activate cytoprotective gene expression ^13^. In cancer cells, loss of KEAP1 function can sustain NRF2 activation, enhancing cystine uptake and GSH biosynthetic capacity ^14^.

Despite substantial research on the cytoprotective functions of GSH, which metabolic processes become limiting as GSH declines remains incompletely understood. Here, using a loss-of-function screen, of cells exposed to GSH synthesis inhibitor buthionine sulfoximine (BSO), a selective inhibitor of glutamate–cysteine ligase ^15^, we report that the cystine reductase TXNDC17 is essential for cell fitness when GSH synthesis and availability are limited *in vitro* and *in vivo*. These findings underscore an important function of GSH in preventing disulfide stress and associated disulfidptosis ^16^. Lastly, we also identified an electrophilic scaffold that induced cell death in GSH-depleted cancer cells that depended on cystine uptake. These findings reveal a metabolic vulnerability that could be pharmacologically exploited to eradicate cancers with high cystine uptake.

## Results

### TXNDC17 protects cancer cells during GSH depletion

To identify factors required for cell survival during GSH depletion, we performed a genome-wide CRISPR screen in KEAP1-mutant H460 lung cancer cells treated with BSO (Fig. 1a). Parallel screens with and without ferroptosis inhibitor liproxstatin-1 (Lip-1) allowed us to distinguish ferroptosis-dependent and independent effects. TXNDC17, a cystine reductase also known as TRP14, emerged as a prominent sensitizing hit in both conditions, whereas loss of the ferroptosis suppressor FSP1 ^17^ sensitized cells only in the absence of Lip-1 (Fig. 1b). We validated these findings using two independent single-guide RNAs targeting TXNDC17, where we could show that loss of TXNDC17 increased BSO sensitivity in A375 and H460 cells, with a corresponding loss of TXNDC17 protein. (Fig. 1c). The same phenotype in A549, H1975 and PC9 cells extended this dependence across different cancer cell lines (Extended Data Fig. 1b–d).

**Fig. 1.**
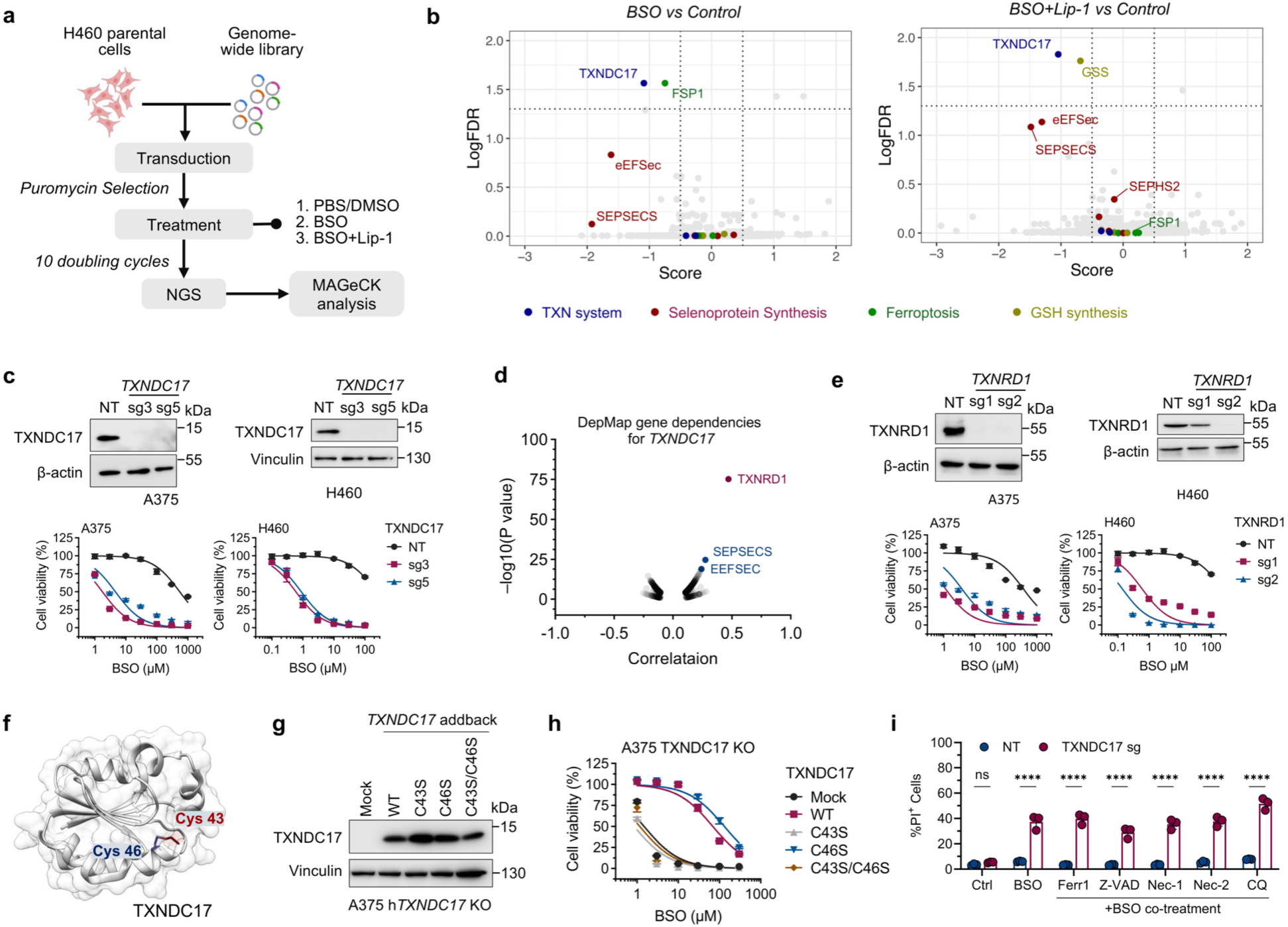
Loss of TXNDC17 sensitizes cells to GSH depletion. **a.** Workflow of the loss-of-function screen using the genome-wide Toronto KnockOut (TKO) CRISPR Library, version 3. **b.** Volcano plots showing gene enrichment in the genetic screen following BSO treatment (left) or combined BSO and Lip-1 treatment (right), with both conditions compared with the vehicle control. **c.** Top: Immunoblots of TXNDC17 in A375 (left) and H460 (right) cells. Bottom: Cell viability following BSO treatment in *TXNDC17*-knockout cells generated using two independent sgRNAs (sg3 and sg5) and the corresponding non-targeting control cells (NT). **d.** DepMap database (version 26Q3) analysis to identify genes co-dependent with *TXNDC17*. **e.** Top: Immunoblots of TXNRD1 in A375 (left) and H460 (right) cells. Bottom: Cell viability following BSO treatment in *TXNRD1*-knockout cells generated using two independent sgRNAs (sg1 and sg2) and the corresponding non-targeting control cells (NT). **f.** Protein structure of TXNDC17 obtained from Protein Data Bank (PDB). PDB ID: 1WOU (pdb_00001wou). The cysteine residues at the catalytic sites are highlighted. **g.** Immunoblot analysis of TXNDC17 expression in A375 *TXNDC17*-knockout cells reconstituted wild-type TXNDC17 or the indicated mutants. **h.** Cell viability following BSO treatment in A375 *TXNDC17*-knockout cells reconstituted wild-type TXNDC17 or the indicated mutants. **i.** Cell death analysis by propidium iodide (PI) staining in H460 *TXNDC17*-knockout cells cultured for 72 h with 100 µM BSO in the presence of the indicated compounds: ferrostatin-1 (Ferr1; 10 µM), Z-VAD-FMK (Z-VAD; 10 µM), necrostatin-1 (Nec-1; 10 µM), necrostatin-2 (Nec-2; 10 µM), or chloroquine (CQ; 25 µM).

Analysis of the Cancer Dependency Map identified thioredoxin reductase 1 (TXNRD1), as the highest co-dependency with TXNDC17 (Fig. 1d). Although TXNRD1 did not score in the primary screen, its loss increased BSO sensitivity in A375 and H460 cells (Fig. 1e), similar to previous reports ^18^. To determine whether TXNDC17-dependent protection required its disulfide exchange activity, we reconstituted *TXNDC17*-knockout A375 cells with wild-type or cysteine-to-serine mutant constructs (Fig. 1f,g). Wild type TXNDC17 restored resistance to BSO, whereas C43S and the C43S/C46S double mutant failed to protect. The C46S variant, however, retained substantial protection (Fig. 1h). The attacking cysteine, Cys43, was therefore essential and sufficient even when the resolving cysteine was absent. These results align with previous enzymatic studies showing that TXNRD1 reduced the mixed disulfide with cystine formed at Cys43 in the C46S variant ^19^.

Motivated by the strong cell death induced by BSO in cells lacking TXNDC17 we next tested whether inhibiting established death pathways could prevent cell death. For this, we treated cells with BSO in combination with a series of cell death inhibitors, namely ferrostatin-1, an inhibitor of ferroptosis; Z-VAD, a pan-caspase inhibitor of caspase-dependent apoptosis; necrostatin-1 or necrostatin-2, inhibitors of necroptosis; or chloroquine, an inhibitor of autophagic flux. Interestingly, none of these treatments prevented cell death (Fig. 1i), indicating that this cytotoxic modality is distinct from the tested forms of cell death.

### GSH depletion exposes a dependence on TXNDC17 for cystine reduction

To better understand the factors regulating cell death upon GSH depletion in *TXNDC17*-deficient cells, we performed a second genome-wide CRISPR screen, this time using *TXNDC17*-deficient H460 cells treated with BSO (Fig. 2a). *SLC7A11* and *SLC3A2*, which encode the xCT transport subunit and its 4F2hc heavy-chain partner ^10^, emerged as prominent resistance hits (Fig. 2b), indicating that the activity of this transporter contributed to the cell death observed in *TXNDC17*-deficient cells treated with BSO. Accordingly, disrupting *SLC7A11* conferred resistance to BSO in *TXNDC17*-deficient cells (Fig. 2c). This genetic rescue placed cystine transport upstream of the toxicity associated with combined loss of TXNDC17 and GSH depletion.

**Fig. 2.**
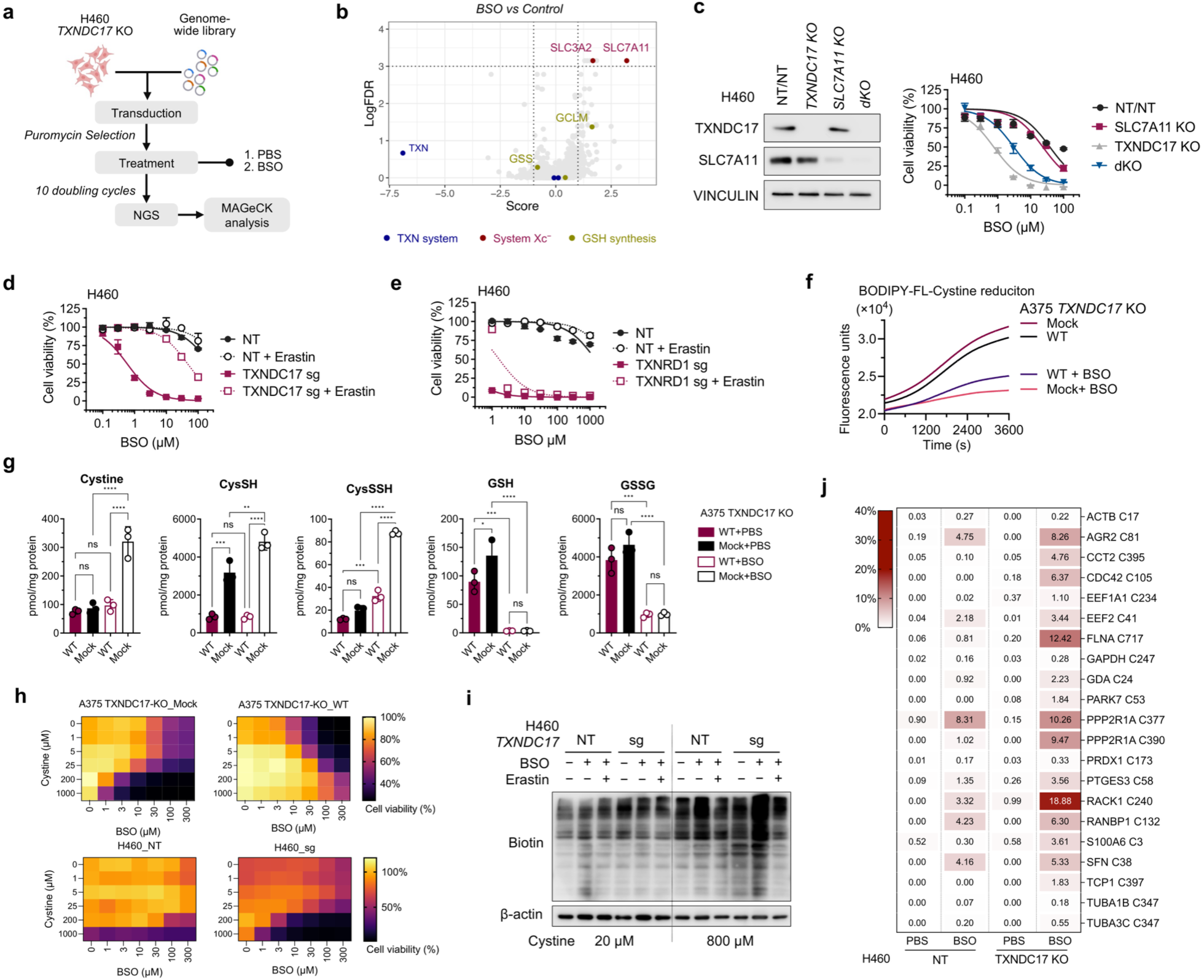
TXNDC17 and GSH cooperate to prevent cystine accumulation and protein cysteinylation. **a.** Workflow of the genome-wide loss-of-function screen using the Toronto KnockOut (TKO) CRISPR Library version 3, to identify regulators of cell death in *TXNDC17*-deficient cells upon BSO treatment. **b.** Volcano plot showing gene enrichment in the genetic screen performed in *TXNDC17*-knockout H460 cells following BSO treatment, compared with the vehicle control. **c.** Left: Immunoblots of TXNDC17 and SLC7A11 in H460 cells of the indicated genotypes. Right: Cell viability following BSO treatment in H460 cells of the indicated genotypes and the corresponding non-targeting control cells (NT/NT). **d.** Cell viability following BSO treatment in H460 *TXNDC17*-knockout cells co-treated with 1 µM erastin or vehicle (DMSO). **e.** Cell viability performed as described in Fig. 2d-using H460 *TXNRD1*-knockout cells. **f.** Kinetic analysis of cystine reduction by TXNDC17, monitored using BODIPY™-FL L-cystine. A375 TXNDC17-knockout cells reconstituted with wild-type TXNDC17 or mock control cells were incubated with 50 µM BSO or vehicle (PBS) for 24 h before the assay. **g.** Quantification of sulfur metabolites by LC–MS/MS. A375 *TXNDC17*-knockout cells reconstituted with wild-type TXNDC17 or mock control cells were incubated with 50 µM BSO or vehicle (PBS) for 24 h. CysSH, cysteine; CysSSH, cysteine persulfide; GSH, reduced glutathione; GSSG, glutathione disulfide. Data are presented as mean ± SD. Statistical analysis was performed using one-way ANOVA followed by Tukey’s multiple-comparisons test. ns, not significant (P > 0.05); *P ≤ 0.05; **P ≤ 0.01; ***P ≤ 0.001; ****P ≤ 0.0001. **h.** Dose-dependent cystine toxicity under conditions of glutathione depletion. Cell viability was assessed in *TXNDC17*-knockout cells reconstituted with wild-type TXNDC17 (top-right) or mock control cells (top-left) and H460 *TXNDC17*-knockout cells (sg; bottom-right) or control cells (NT; bottom-left) following BSO treatment in medium containing different concentrations of L-cystine. **i.** Analysis of protein cysteinylation in H460 *TXNDC17*-knockout cells. Cells were incubated for 24 h in medium containing 20 µM or 800 µM cystine and treated with PBS or 50 µM BSO in combination with 1 µM erastin or vehicle (DMSO), as indicated. **j.** Site-resolved protein S-cysteinylation analysis in H460 *TXNDC17*-targeted and NT control cells treated with 50 µM BSO or PBS for 24 h in the presence of 500 nM Lip-1. The heatmap displays S-cysteinylation site occupancy values from an initial the primary DIA analysis based on a DDA derived spectral library.

Similarly, the system x_c_^−^ inhibitor erastin fully rescued BSO-induced viability loss in *TXNDC17*-deficient H460 cells and protected *TXNDC17*-knockout A375 cells reconstituted with an empty vector or inactive mutant (C43S) (Fig. 2d and Extended Data Fig. 2a). This protection extended across multiple cell models (Extended Data Fig. 2b). Notably, erastin rescue distinguished TXNDC17 deficiency from TXNRD1 loss: although both perturbations sensitized cells to BSO, erastin failed to rescue *TXNRD1*-deficient H460 or A375 cells (Fig. 2e and Extended Data Fig. 2c). These results confirm that TXNRD1 has more functions than cystine reduction via TXNDC17 as restricting cystine uptake selectively relieved the vulnerability exposed by TXNDC17 loss. This mechanistic distinction suggests that the well-established synergy between TXNRD1 loss and GSH depletion ^18,20^ reflects broader redox disruption that drives cell death.

These rescue experiments prompted us to examine cellular cystine-reducing capacity using a fluorogenic BODIPY–cystine probe ^21^. Unexpectedly, and unlike previously reported ^21^, *TXNDC17*-deficient cells showed greater probe reduction than TXNDC17-proficient cells (Fig. 2f). Cells lacking TXNDC17 were characterized by higher NRF2, xCT and TXNRD1 protein signals and GSH levels (Fig. 2g and Extended Data Fig. 2d), raising the possibility that an adaptive response preserves disulfide reduction following TXNDC17 loss. In line with a direct contribution from GSH, BSO suppressed probe reduction to a much greater extent in TXNDC17-deficient cells (Fig. 2f and Extended Data Fig. 2e), highlighting an important and direct contribution of GSH to the reduction of the disulfide probe.

This observation suggests that cystine reduction capacity is shared between TXNDC17 and GSH. LC–MS/MS profiling of sulfur metabolites revealed how this conditional dependency was associated with changes in cystine handling. Under basal conditions, A375 *TXNDC17*-deficient cells contained more GSH and free cysteine than *TXNDC17*-proficient cells, whereas cystine abundance was comparable. BSO markedly depleted GSH and lowered GSSG in both groups. Despite this shared depletion, cystine accumulated predominantly in *TXNDC17*-deficient cells, while cells expressing wild-type TXNDC17 limited its accumulation (Fig. 2g). Free cysteine remained higher in mock cells before and after BSO treatment. The combined perturbation therefore led to cystine accumulation without exhausting the bulk free cysteine pool. The profiling further revealed opposing changes in persulfide pools: BSO increased cysteine persulfide (CysSSH), most prominently in *TXNDC17*-deficient cells, while lowering glutathione persulfide (GSSH) in both groups (Fig. 2g and Extended Data Fig. 3a).

In accordance with a critical contribution of cystine to the toxic effect observed upon TXNDC17 loss, higher extracellular cystine markedly increased BSO sensitivity in cells lacking TXNDC17 (Fig. 2h). Similarly, disrupting NRF2 partially attenuated the loss of viability (Extended Data Fig. 2f), while activating NRF2 with KEAP1/NRF2 interaction inhibitor KI696 sensitized H1975 and PC9 cells, two KEAP1-wild-type lung cancer lines, under high-cystine conditions (Extended Data Fig. 3b).

We next examined whether impaired cystine handling was accompanied by increased protein S-cysteinylation. DIA proteomics identified 21 sites, with apparent occupancy estimated as modified/total peptide precursor signal. Twelve sites showed a greater BSO response in *TXNDC17*-knockout H460 cells compared to controls, including FLNA Cys717, CCT2 Cys395 and RACK1 Cys240 (genotype-by-treatment interaction, adjusted P < 0.05). PRDX1 Cys173 occupancy increased in both genotypes (Fig. 2j). Extended-library analysis additionally identified greater BSO responses at PRDX6 Cys47 and Cys91 in knockout cells, whereas Cofilin-1 Cys139 occupancy increased upon BSO treatment within the knockout genotype (Extended Data Fig. 3d). These findings connect impaired cystine handling to site-specific mixed-disulfide modifications in essential proteins involved in cytoskeletal organization and protein homeostasis, resembling features of disulfide stress-induced disulfidptosis. Disulfidptosis is a cell death modality associated with excess intracellular disulfide stress in cells with high xCT activity, however classically triggered by glucose starvation ^16^.

Further characterization of the transcriptomic response of BSO-induced cell death in TXNDC17-deficient cells identified a marked p53 response that was attenuated by erastin or wild-type TXNDC17 reconstitution. However, the genetic deletion of p53 did not rescue viability (Extended Data Fig. 4a–c).

Altogether, these results establish that cystine availability and uptake are key determinants of cell death induced by combined TXNDC17 loss and GSH synthesis inhibition through a mechanism involving proteomic cysteinylation and oligomerization.

### GSH depletion suppresses TXNDC17-deficient xenograft growth

Having established the mechanistic basis of GSH-depletion-induced cell death in *TXNDC17*-deficient cells, we next assessed whether these effects are also observed in vivo. Female athymic nude mice bearing subcutaneous control or *TXNDC17*-deficient H460 xenografts received subcutaneous BSO or vehicle every other day once tumors reached 50–100 mm^3^ (Fig. 3a). BSO markedly suppressed the growth of *TXNDC17*-deficient xenografts over 18 days, with a smaller response in control tumors. Endpoint tumor weights followed the same pattern, with the lowest weights observed in BSO-treated *TXNDC17*-deficient xenografts, while host body weights remained similar across groups (Fig. 3b–d). The genotype-dependent response therefore extended the vulnerability identified in cultured cells to an established xenograft model. The increased cysteinylation of cytoskeletal proteins in cultured cells prompted us to ask whether cytoskeletal disulfide bonding accompanied this response in tumors. We therefore examined disulfide bonding of actin and drebrin in tumor lysates as markers of disulfidptosis. Higher-molecular-weight species, most prominently actin, accumulated in BSO-treated *TXNDC17*-deficient tumors and were attenuated under reducing conditions (Fig. 3e). Such reduction-sensitive changes in the electrophoretic migration of actin cytoskeleton proteins are characteristic biochemical features of disulfidptosis, in which aberrant disulfide bonding compromises cytoskeletal integrity ^16^.

**Fig. 3.**
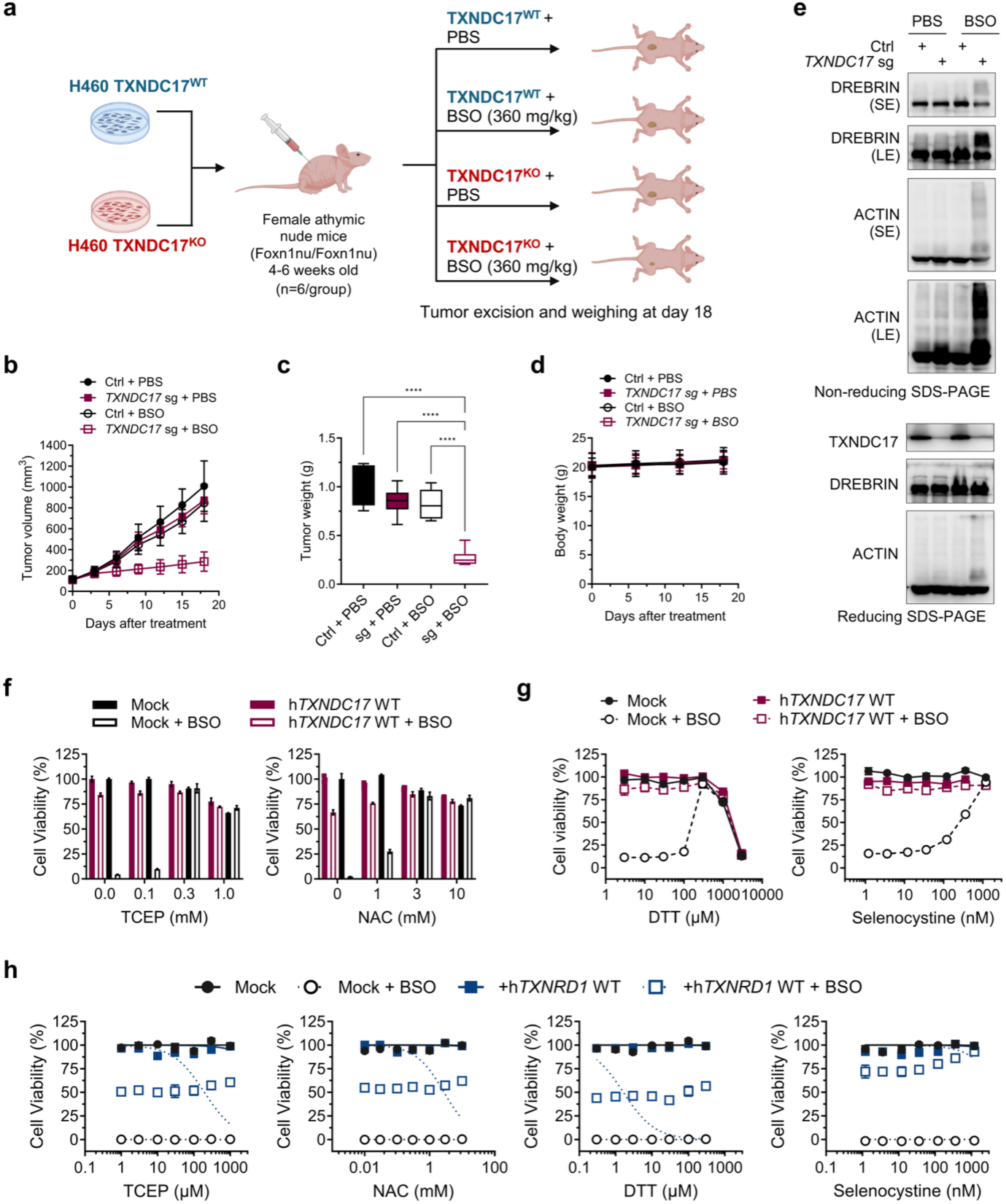
GSH depletion suppresses the growth of TXNDC17-deficient xenografts and induces disulfidptosis. **a.** Workflow of the tumor xenograft experiment. **b.** Tumor volume over time in athymic nude mice subcutaneously inoculated with H460 *TXNDC17*-knockout cells or the corresponding control cells. Once tumors reached 50–100 mm^3^, mice received subcutaneous injections of BSO (360 mg kg^−1^ in 100 µL PBS) or vehicle alone every 2 days. **c.** Tumor weights at the experimental endpoint corresponding to the tumors shown in Fig. 3a. **d.** Body weights of the host nude mice over the course of the experiment shown in Fig. 3a. **e.** Immunoblot analysis of dissected tumor tissues to assess markers of disulfide stress using non-reducing (top) and reducing (bottom) gels. SE, short exposure; LE, long exposure. **f.** Cell viability following treatment with tris(2-carboxyethyl)phosphine (TCEP; left) or N-Acetyl-L-Cystein (NAC; right) under conditions of glutathione depletion induced by 50 µM BSO in A375 *TXNDC17*-knockout cells reconstituted with wild-type TXNDC17 or the corresponding mock control cells. **g**. Cell viability following treatment with dithiothreitol (DTT; left) or seleno-L-cystine (selenocystine; right) assessed as described in Fig. 3f in A375 *TXNDC17*-knockout cells reconstituted with wild-type TXNDC17 or the corresponding mock control cells. **h.** Cell viability assessed as described in Fig. 3f using A375 *TXNRD1*-knockout cells expressing wild-type *TXNRD1* or the corresponding mock control cells.

To assess whether redox-modulating interventions could alleviate this vulnerability, we treated BSO-exposed A375 cells with disulfide reductants or sulfur- and selenium-containing supplements. TCEP, DTT, selenocystine, and N-acetyl-L-cysteine restored the viability of *TXNDC17*-deficient cells treated with BSO (Fig. 3f,g). Of note, the same reductants were unable to rescue *TXNRD1*-deficient A375 cells exposed to BSO (Fig. 3h). These rescue experiments further distinguish the consequences of GSH depletion in combination with one of the two reductases and are consistent with TXNRD1 having redox functions beyond supplying reducing equivalents to TXNDC17.

### Electrophilic fragments enhance cystine-dependent toxicity

Finally, in order to establish pharmacological tools that could recapitulate our observations, we screened 960 electrophilic compounds in BSO-pretreated H460 cells cultured in 800 μM cystine and Lip-1. We used erastin as a counter-screen to identify compounds that reduced the viability signal and showed relative rescue when cystine transport was inhibited (Fig. 4a,b). We profiled six candidate fragments in which BSO enhanced viability loss, and erastin or *SLC7A11* disruption attenuated the response; we called these cystine toxicity enhancers (CTE) (Fig. 4c,d and Extended Data Fig. 5a). CTE-2 stood out for its greatest activity during GSH depletion and the protection conferred by both interventions. It induced dose-dependent viability loss in BSO-treated A375, H460, A549, H1975, H1299 and EKVX cells, with comparatively limited effects without BSO. Erastin consistently suppressed the cell death induced by the combination of BSO and CTE-2 (Fig. 4e). Disrupting GSH synthesis enzymes GCLC or GCLM also markedly sensitized cells to CTE-2, with protection by erastin (Extended Data Fig. 5b).

**Fig. 4.**
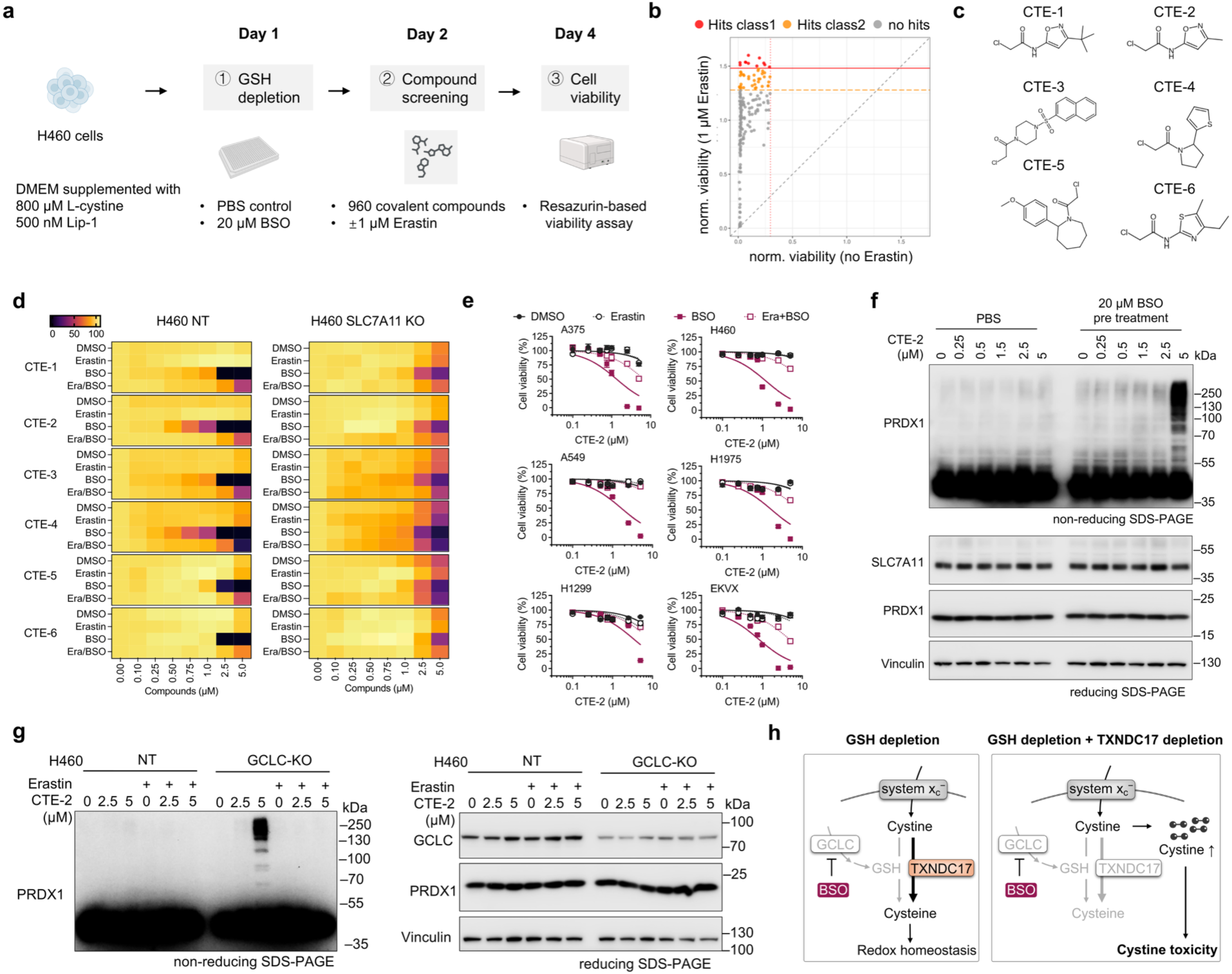
Electrophilic fragments induce cystine-dependent disulfidptosis in GSH-depleted cells. **a.** Workflow of the screen using a library of electrophilic covalent probes. **b.** Summary of the primary screen performed using each library compound at 5 µM. The X-axis represents cell viability in the absence of 1 µM erastin, whereas the Y-axis represents cell viability in the presence of 1 µM erastin. Cell viability values were normalized to those of DMSO-treated control wells. Compounds showing cytotoxicity in the absence of erastin and an erastin-mediated rescue of compound-induced cytotoxicity exceeding the indicated cutoff values (hit classes 1 and 2) were called cystine-toxicity enhancers (CTE) and were selected for further hit validation. **c.** Chemical structures of the CTEs identified through hit validation. **d.** Cell viability following treatment with the CTEs depicted in Fig. 4c. H460 *SLC7A11*-knockout cells and the corresponding non-targeting control cells (NT) were incubated with 20 µM BSO or vehicle (PBS) for 24 h, followed by treatment with the indicated concentrations of each CTE for an additional 48 h. **e.** Cell viability performed as described in Fig. 4d using CTE-2 in the indicated cancer cell lines. **f.** Immunoblot analysis of PRDX1 oligomerization. Parental H460 cells were incubated with 20 µM BSO or vehicle (PBS) for 24 h, followed by treatment with the indicated concentrations of CTE-2 for an additional 24 h. PRDX1 was detected using non reducing (top) and reducing (bottom) gels. **g.** Immunoblot analysis of PRDX1 oligomerization. H460 GCLC-knockout cells and the corresponding non-targeting control cells (NT) were incubated with the indicated concentrations of CTE-2 for an additional 24 h. PRDX1 was detected using non reducing (left) and reducing (right) gels. **h**. Proposed model for cystine toxicity during combined GSH depletion and TXNDC17 loss. BSO inhibits the GCLC-catalyzed step of GSH synthesis. During GSH depletion (left), TXNDC17 sustains the reduction of imported cystine to cysteine, helping maintain redox homeostasis. When TXNDC17 is deficient (right), continued cystine uptake through system x_c_⁻ exceeds the cell’s capacity to reduce cystine. The resulting intracellular cystine accumulation promotes disulfidptosis.

Given the increased S-cysteinylation of PRDX1 identified above, we asked whether CTE-2 could alter the migration of this redox-sensitive protein. CTE-2 increased higher-molecular-weight PRDX1-reactive species on non-reducing immunoblots after BSO pretreatment, most prominently at 5 μM (Fig. 4f). These species also increased in GCLC-knockout cells and were attenuated by erastin (Fig. 4g). This pattern is consistent with protein thiol–disulfide imbalance, providing a biochemical signature compatible with the disulfide stress characteristic of disulfidptosis. These data identify CTE-2 as a phenotypic lead that, in combination with BSO, can directly induce disulfidptosis; however, the direct target of CTE-2 remains to be established.

## Discussion

Our findings reveal a conditional requirement for TXNDC17 in cancer cells that emerges upon GSH depletion. TXNDC17 was previously established as cystine-reductase and protein-decysteinylase ^19,21,22^. Furthermore, we show that GSH-dependent processes compensate for TXNDC17 loss enough to sustain cell fitness; however, combined TXNDC17 loss and GSH depletion cause cystine accumulation and cystine-dependent cell death. Notably, cystine accumulates despite an increase in the bulk free cysteine pool, arguing against simple cellular cysteine depletion as the cause of toxicity. These findings highlight a role for cystine reduction beyond cysteine supply: cells must also manage the disulfide load imposed by continued cystine uptake.

The dependence on cystine uptake, genetic and pharmacological rescue experiments, and reduction-sensitive cytoskeletal species support the occurrence of disulfidptosis following combined TXNDC17 loss and GSH depletion. Previous work established that glucose deprivation can drive disulfidptosis by limiting NADPH availability and promoting aberrant disulfide bonding in cytoskeletal proteins ^16^. Our findings identify a vulnerability in the machinery that processes imported cystine, exposed without forced glucose restriction. Increased protein S-cysteinylation further connects impaired cystine metabolism to protein thiol modification. Given that TXNDC17 can directly reverse PRDX2 S-cysteinylation in vitro, accumulation of these mixed disulfides could reflect both increased formation as cystine accumulates and impaired removal following loss of TXNDC17 and GSH-dependent reducing capacity ^21^. The relative contributions of cystine reduction and direct protein decysteinylation to TXNDC17-mediated protection, and the modifications responsible for cytoskeletal dysfunction, remain to be established. Interestingly, we also observed a strong stabilization of p53 and the associated transcriptional signature in cells undergoing disulfide stress. Although genetic deletion of p53 did not rescue viability, this does not exclude a homeostatic role for p53 in limiting cystine influx. p53 can repress *SLC7A11* expression and reduce cystine uptake ^23^. We therefore speculate that p53 activation could provide negative feedback by restricting cystine entry when intracellular reductive capacity is compromised. Such feedback may be overwhelmed by combined TXNDC17 loss and pronounced GSH depletion, yet it may influence the toxicity threshold under milder conditions, such as partial inhibition of GSH synthesis and/or TXNDC17 activity.

The preservation of bulk cysteine also points to the flexibility of cellular sulfur metabolism. Transsulfuration can compensate for TXNDC17 loss in some nutrient and tissue contexts, while cystine cleavage provides an alternative source of cysteine under disulfide reductase deficiency ^21,24^. These pathways could help maintain cysteine availability without fully preventing cystine accumulation in our models.

Comparing TXNDC17 and TXNRD1 deficiency helps define the specificity of this dependency. Although both perturbations sensitized cells to GSH depletion, restricting cystine uptake selectively rescued TXNDC17-deficient cells, and the tested reductant interventions failed to rescue TXNRD1-deficient cells. These differences suggest that cystine metabolism is a dominant vulnerability following TXNDC17 loss, whereas TXNRD1 deficiency compromises additional processes that remain essential when cystine uptake is restricted. This distinction could be context-dependent as TXNRD1 inhibition was shown to sensitize glucose-starved glioblastoma cells to disulfidptosis ^25^. Thus, shared sensitivity to GSH depletion can arise from different limiting functions within the thioredoxin system. Importantly, the xenograft response supports the relevance of this dependency *in vivo*, while the cystine titrations indicate that extracellular nutrient availability can influence its magnitude.

As a first step toward providing tools to trigger this form of cell death, CTE-2 offers a chemical starting point for exploiting the vulnerability revealed by TXNDC17 loss. Its enhancement of cystine-dependent toxicity during GSH depletion, together with the accumulation of PRDX1 high-molecular-weight species, reproduces several features of the genetic phenotype. This convergence suggests that electrophilic compounds can perturb processes required to tolerate cystine uptake when GSH availability is limited. However, these phenotypic similarities do not establish TXNDC17 as the molecular target of CTE-2. Defining its direct targets and their contribution to cell death will be essential to determine how the compound disrupts cystine handling and to develop more selective chemical probes.

Altogether, our findings identify TXNDC17 and GSH as a critical defense against cystine toxicity and establish a critical regulatory node in disulfidptosis. Targeting the capacity to process imported cystine may therefore convert high cystine uptake from a survival advantage into a therapeutic liability by directly triggering this form of cell death.

## MATERIALS AND METHODS

### Chemicals

L-Buthionine-sulfoximine (BSO; cat. no. B25159), erastin (329600), protamine sulfate (P3369), L-cystine (V8755), seleno-L-cystine (545996), tris (2-carboxyethyl) phosphine hydrochloride (TCEP-HCl; C4706), and liproxstatin-1 (SML1414), were purchased from Sigma-Aldrich. N-ethylmaleimide (NEM; HY-D0843) and KI696 (HY-101140) were obtained from MedChemExpress. The compounds identified from the electrophilic covalent probe library (Z241744828, Z90122381, Z199507050, Z56896883, Z147646956, Z561062354, Z425859482) were purchased.

### Cell culture

The human cancer cell lines A375 (CRL-1619), H1975 (CRL-5908), A549 (CRM-CCL-185), and H460 (HTB-177) were obtained from the American Type Culture Collection (ATCC). PC-9 (90071810) cells were purchased from Sigma-Aldrich. EKVX (CVCL_1195) was a kind gift from Almut Schulze. Cells were cultured in high-glucose DMEM GlutaMAX (Gibco, 31966-021) or RPMI 1640 medium (Gibco, 11875093; for PC-9 cells), supplemented with 10% fetal bovine serum (FBS; Gibco, A5256701) and 1% penicillin-streptomycin (Gibco, 15140122). Cells were maintained at 37°C in a humidified atmosphere containing 5% CO_2_ and were routinely tested for mycoplasma contamination. Unless otherwise indicated, cells were maintained in glucose-replete culture medium without additional selenium supplementation.

### Molecular cloning

sgRNAs were designed using the VBC score (https://www.vbc-score.org/). Annealed oligonucleotides (synthesized by Microsynth) were ligated into BsmBI (NEB, #R0739)-digested lentiCRISPRv2-puro, lentiCRISPRv2-blast, or lentiCRISPRv2-hygro vectors (Addgene, #98290, #98293, and #98291, respectively) with T4 ligase (NEB, M020L). All sgRNA sequences used for the study were listed in Extended Data Table 1.

cDNA fragments for codon-optimized TXNDC17 wildtype, C43S and C46S mutants were obtained from Integrated DNA Technologies (IDT). These cDNAs were cloned into linearized lentiviral vector p442-IRES (kindly provided by Prof. Dr. Marcus Conrad) using Clone Express MultiS One Step Cloning Kit (Vazyme, #VB-C113-02).

sgRNA and cDNA insertion into the respective plasmids was confirmed by Sanger sequencing (Microsynth).

### Lentivirus production and transduction

For lentiviral particle production, human embryonic kidney (HEK) 293T cells were transfected with the packaging plasmids pMDLg/pRRE (Addgene, #12251) and pRSV-Rev (Addgene, #12253), together with the envelope plasmid pCMV-VSV-G (Addgene, #8454), using X-tremeGENE HP DNA Transfection Reagent (Sigma-Aldrich, 6366546001). After 48 hours, the supernatant containing lentiviral particles was collected, filtered through a 0.44-μm-pore membrane, and used for transduction in the presence of 8 µg/mL protamine sulfate. Transduced cells were selected with the corresponding antibiotics, and gene knockout was validated by loss of protein expression by immunoblotting.

### Generation of clonal H460 TXNDC17^KO^ by transient plasmid transfection

H460 parental cells were transiently transfected with a lentiCRISPRv2-puro plasmid containing a sgRNA targeting *TXNDC17* (sg3; Extended Data Table 1), using the transfection reagent described above. Transfected cells were selected with 1 μg/mL puromycin, and surviving cells were expanded and subjected to single-cell cloning. Individual clones were cultured in medium with or without 30 μM BSO, and clones that failed to survive BSO treatment were selected, and loss of TXNDC17 protein expression was confirmed by immunoblotting. The one validated *TXNDC17*-knockout clone (Cl.4) was subsequently used for the CRISPR screen.

### Genome-wide CRISPR-Cas9 knockout screens

H460 parental and H460 transient-*TXNDC17*^KO^ Cl.4 cells were transduced with a human genome-wide knockout library, Toronto KnockOut (TKO) CRISPR Library Version 3 (Addgene, #90294). Transduced cells were maintained in the presence of 1 μg/mL puromycin (InvivoGen, ant-pr-1) and subsequently treated with 50 μM BSO in the presence or absence of 500 nM Lip-1 for 72 hours. The surviving populations were maintained until they reached 10 doublings, after which cells were harvested and snap-frozen in liquid nitrogen. Genomic DNA was extracted using the QIAamp DNA Blood Maxi Kit (QIAGEN, 51194). The sgRNA regions of the library plasmids were amplified using a mixture of forward primers containing stagger sequences to increase base diversity and reverse primers containing distinct barcode sequences (Extended Data Tables 2 and 3). PCR was performed using Phanta Max Master Mix (Absource, P515-03) under the following conditions: initial denaturation at 95°C for 3 min, followed by 28 cycles of denaturation at 95°C for 30 s, annealing at 54°C for 30 s, and extension at 72°C for 30 s. A final extension was performed at 72°C for 5 min. The amplicons were purified using the Wizard SV Gel and PCR Clean-Up System (Promega, A9282) and sequenced on a NextSeq 2000 system (Illumina) using 100-bp single-end reads, with a sequencing depth of approximately 40 million reads per sample. 500-fold library coverage was maintained throughout the process. Raw read counts were analyzed using the MAGeCK pipeline ^26^, and gene enrichment relative to the corresponding controls was visualized with RStudio.

### Immunoblotting

Cells were washed with ice-cold PBS twice and lysed in RIPA buffer [20 mM Tris-HCl, 150 mM NaCl, 1 mM EDTA, 1 mM EGTA, 1% NP-40, 1% deoxycholate] supplemented with protease inhibitor cocktail (Roche, 04693159001). For assessment of PRDX1 oligomerization, N-ethylmaleimide (NEM) was added to the RIPA buffer at a final concentration of 10 mM. After centrifuging the lysate for 30 min at 4°C and 18000g, supernatants were collected. Protein concentrations were measured using Pierce BCA Protein Assay Kit (Thermo Scientific, 23227) with bovine serum albumin (BSA) as protein standard. 12% SDS-polyacrylamide gel electrophoresis was used to separate proteins followed by electro-transfer to polyvinylidene difluoride (PVDF) membranes. Membranes were blocked in 5% non-fat milk or 5% BSA in Tris-buffered saline with 0.1 % Tween 20 (TBS-T) for 1 h at room temperature. Primary antibodies were diluted according to the manufacturer’s suggestion in a 5% BSA or 5% non-fat milk in TBS-T solution and incubated with the membrane overnight at 4°C. All antibodies used in this study are listed in Extended Data Table 4. After 3 times washes in TBS-T, membranes were incubated with 1:3000 dilution of the secondary antibodies (Cell Signalling Technology, anti-rabbit IgG-HRP linked (#7074), anti-mouse IgG-HRP linked (#7076), or anti-rat IgG-HRP linked (#7077) at RT for 1 hour. Proteins were detected using Immobilon Western chemiluminescence HRP-Substrate (Millipore, WBKLS0050). Images were acquired using detection systems Amersham ImageQuant 800 (Cytiva).

### Cell viability assays

Cell viability was determined based on cellular reducing capacity using a resazurin-based assay. Cells were seeded in a 96-well plate at a density of 2,000–3,000 cells/well and treated with the designated compounds. Following 48–72 hours of incubation, the medium was replaced with resazurin-containing DMEM (diluted 1:150 from a 0.5% stock solution in medium) and incubated for an additional 4 hours. Fluorescence was measured on a Spark® microplate reader (Tecan) using a 540±35 nm excitation filter and a 590±20 nm emission filter. Raw fluorescence values were normalized to those of the corresponding vehicle-treated control wells, which were set to 100%.

### Cellular BODIPY FL L-cystine reduction assay

Cellular cystine-reducing capacity was assessed using BODIPY FL L-cystine according to a published procedure with modifications^4^. Cells were seeded in lumox® 96-well plate at 3,000 cells per well and cultured in DMEM 10% FBS for 4 hours. Cells were exposed to BSO or vehicle (PBS) for the next 24h. Then, BODIPY FL L-cystine (MCE; HY-D1605) was added at a final concentration of 1 μM in 100 μl of PBS plus GlutaMAX™.

Fluorescence was recorded using a TECAN plate reader (TECAN, Switzerland) at 1-minute intervals for at least 30 minutes at 37°C, with excitation at 505 nm and emission at 520 nm.

### LC-MS/MS analysis of sulfur metabolites

TXNDC17-knockout cells carrying wildtype TXNDC17 or mock control cells were seeded at 0.35 × 10^6^ cells per well in 6-well plates and cultured in medium containing vehicle (PBS) or 50 μM BSO for 24 hours. Cells were then washed twice with ice-cold PBS. The bottom of each 6-well plate was snap-frozen in liquid nitrogen and stored at −80°C until further processing.

For metabolite extraction, the plates were thawed, and 500 μL per well of extraction buffer [5 mM HPE-IAM in 70% methanol and 50 mM sodium acetate (pH 6.5)] was added to lyse the cells. The lysates were collected into tubes by scraping, disrupted by probe sonication at 30% intensity for 10 s twice, and incubated at 37°C for 20 min. The lysates were then centrifuged at 10,000g for 10 min at 4°C. A 50 μL aliquot of each supernatant was collected and mixed with 100 μL of 0.1% formic acid. The protein pellets were air-dried, dissolved in 300 μL of 0.2 M NaOH solution containing 0.1% SDS, and incubated at 60°C for 30 min for BCA-based protein normalization. For analysis, 10 μL of the samples diluted with 0.1% formic acid was mixed with 10 μL of 50 nM isotope-labeled internal standards, and 10 μL of the final mixture was injected for LC-MS/MS analysis.

Analyses were performed by LC-ESI-MS/MS using an LCMS-8060RX mass spectrometer (Shimadzu) coupled to a Nexera UHPLC system (Shimadzu). Metabolites were separated on a YMC-Triart C18 column (50 × 2.0 mm inner diameter) using mobile phase A (0.1% formic acid in water) and mobile phase B (0.1% formic acid in methanol), with a linear gradient from 3% to 90% B over 15 min, at a flow rate of 0.2 mL/min and a column temperature of 40 °C. The ESI probe, desolvation line, and heat block temperatures were set to 300, 250, and 400 °C, respectively, and the nebulizer, heating, and drying nitrogen gas flow rates were set to 3, 10, and 10 L/min, respectively. Metabolites were identified and quantified by multiple reaction monitoring (MRM). MRM parameters for each metabolite and the synthesis of isotope-labeled internal standards were described previously ^27,28^.

### Protein cysteinylation analysis

Protein cysteinylation was assessed by labeling cells with biotinylated cysteine followed by western blotting, using an adapted published protocol^4^. Cells were seeded in 100 mm dishes at 2.0 x 10^6^ cells per dish and cultured for 4 hours. Cells were exposed to BSO and/or erastin for 24h in regular DMEM medium. N-Biotinyl-L-cysteine (MCE; HY-128466) was added at 250 μM in DMEM cystine and FBS free medium for 1 hour at 37°C.

The labeling medium was removed, cells were washed 2 times with ice-cold PBS, and samples were collected in 150 μl of lysis buffer containing 50 mM Tris-HCl (pH 7.5), 2 mM EDTA, 150 mM NaCl, 1% Triton, 10 mM N-ethylmaleimide (NEM), and protease inhibitor cocktail. Cells were lysed by three freeze-thaw cycles. Lysates were centrifuged at 1370 x g for 10 minutes at 4°C and the supernatants were collected for immunoblot analysis.

### Sample preparation for protein bottom-up mass spectrometric analysis

TXNDC17-knockout and H460 cells were cultured in 10-cm dishes were treated with or without BSO (50 μM) in the presence of Lip1 (500 nM) for 24 h (n=5 per group). After treatment, dishes were washed twice with 15 mL ice-cold, degassed PBS. Proteins were precipitated by adding 1 mL of degassed 20% (w/v) trichloroacetic acid (TCA) to fully cover the cells, followed by incubation at 4 °C for 30 min. Cells were scraped and centrifuged at 13,000 × g for 10 min at 4 °C to collect protein pellets. Pellets were washed with 1 mL of 10% (w/v) TCA and centrifuged at 13,000 × g for 10 min at 4 °C without disturbing the pellet. The washing step was repeated once with 1 mL of 5% (w/v) TCA. After centrifugation, the pellets were dissolved in 400 µL of degassed lysis buffer (8 M urea, 1% SDS, 250 mM HEPES, 100 mM NEM in water, pH 7.5). Samples were incubated in the dark at 37 °C for 1 h with shaking. Proteins were then reprecipitated by adding 1.6 mL of ice-cold acetone, vortexing, and incubating at −20 °C overnight. Samples were subsequently transferred to −80 °C until further processing.

Precipitates were thawed on ice and pelleted (13,000 × g, 10 min, 4 °C), washed with 1 ml ice-cold 70% acetone, briefly air-dried and dissolved in 200 µl 1% SDS, 1% Triton X-100 in 2-(N-morpholino)ethanesulfonic acid buffer pH 6.7 by probe sonication (about 1 min). After centrifugation (13,000 × g, 5 min, 11 °C) the supernatant was transferred to a protein low-bind tube and the protein concentration determined by bicinchoninic acid assay.

For each sample 10 µg protein in 20 µl buffer (0.5 mg/ml) were alkylated without prior reduction with 50 mM iodoacetamide (20 min, room temperature, dark). Proteins were processed by protein aggregation capture ^29^: 5 µl of a 20 mg/ml 1:1 mix of hydrophilic and hydrophobic carboxylate-modified magnetic beads (beads to protein 10:1) were added, ethanol was added to 50% (v/v), and samples were incubated for 15 min at 24 °C and 1000 rpm. Beads were collected on a magnetic rack, washed three times with 200 µl 80% ethanol and once with 200 µl acetonitrile, and resuspended in 20 µl 50 mM triethylammonium bicarbonate with 0.2 µg trypsin. Digestion was carried out overnight (13-16 h) at 37 °C with interval mixing at 1000 rpm, followed by addition of another 0.2 µg trypsin and 3 h at 37 °C.

For spectral library generation, half of the digests of the five *TXNDC17* knockout samples treated with buthionine sulfoximine (BSO) were pooled, dried, dissolved in 0.1% trifluoroacetic acid and fractionated with BioSPE PepFrac 200 µl tips (Affinisep, protocol of manufacturer); the fractions were measured in eight data-dependent acquisition runs.

### Mass spectrometric analysis of peptides from cell lysates

Peptides (amount corresponds to 0.8 µg of the initial sample) were loaded on Evotips and separated on an Evosep One using the Whisper Zoom 20 samples-per-day method on an Aurora Elite C18 UHPLC column (15 cm × 75 µm, CSI; IonOpticks). The column was coupled via a CaptiveSpray ion source to a timsTOF Pro 2 mass spectrometer (Bruker Daltonics). For both data-dependent and data-independent acquisition modes, the trapped ion mobility spectrometry accumulation and ramp times were set to 100 ms (100% duty cycle, 9.43 Hz), and MS spectra were recorded from m/z 100 to 1,700. Collision energy was ramped linearly as a function of ion mobility, from 20 eV at 1/K0 = 0.60 V·s/cm^2^ to 59 eV at 1/K0 = 1.60 V·s/cm^2^.

Library samples were acquired in data-dependent acquisition parallel accumulation–serial fragmentation (ddaPASEF) mode across an ion mobility range of 0.70–1.50 V·s/cm^2^. Each 1.17 s cycle consisted of one MS1 survey scan followed by 10 PASEF MS/MS ramps. Precursors with charge states up to 5 were selected using a polygon filter in the m/z–ion mobility plane. Precursors above an intensity threshold of 1,200 were fragmented until they reached a target intensity of 14,500, and were then dynamically excluded for 0.4 min. The quadrupole isolation width was 2 m/z at m/z 700 and 3 m/z at m/z 800.

Samples were acquired in data-independent acquisition PASEF (diaPASEF) mode across an ion mobility range of 0.65–1.45 V·s/cm^2^. Each cycle consisted of one MS1 scan followed by 8 diaPASEF ramps, with an estimated cycle time of 0.95 s. Each ramp contained 3 isolation windows, giving a total of 24 non-overlapping windows of 25 m/z width that covered m/z 400–1,000. The ion mobility boundaries of the windows were adjusted along the peptide precursor distribution in the m/z–1/K0 plane, within 1/K0 0.64–1.37 V·s/cm^2^ (Extended Data Table 5). Each acquisition cycle consisted of one MS1 scan followed by 8 data independent aquision PASEF ramps, each isolating 3 windows (24 windows in total, 25 m/z each, no overlap). For each window, the m/z range and ion mobility range (1/K0, V·s/cm^2^) are given. Collision energy was ramped linearly with ion mobility from 20 eV at 1/K0 0.60 to 59 eV at 1/K0 1.60 V·s/cm^2^.

### Mass spectrometric data analysis

An initial spectral library was generated from eight ddaPASEF runs in Spectronaut 19.8 (Biognosys) using the Pulsar search engine against the human UniProt reference proteome (UP000005640, 83,385 entries, downloaded 16 April 2025), the same database as used in DIA-NN. Trypsin/P was set as a specific enzyme, allowing up to two missed cleavages and peptide lengths of 7–52 amino acids. No fixed modifications were set. Variable modifications (up to five per peptide) were carbamidomethylation, N-ethylmaleimide, hydrolyzed N-ethylmaleimide and cysteinylation of cysteine, oxidation of methionine and protein N-terminal acetylation. MS1 and MS2 tolerances were determined dynamically, and the false discovery rate was set to 1% at peptide-spectrum match, peptide and protein level. The library contained the 3–6 most intense b and y fragment ions per precursor (m/z 200–3,000), and indexed retention time values were calibrated by deep-learning-assisted regression.

To enable protein and peptide identification and quantification, the 20 diaPASEF runs were analyzed with DIA-NN 2.7.0 ^30^ in library-based mode, using the ddaPASEF spectral library together with the human UniProt reference proteome (UP000005640, 83,385 entries, downloaded 16 April 2025), the MaxQuant contaminant database and indexed retention time peptide sequences. Trypsin was set as the protease, allowing one missed cleavage, peptide lengths of 7–30 amino acids, precursor charges of 1–4 and N-terminal methionine excision. Variable modifications (up to three per peptide) were oxidation of methionine and carbamidomethylation, N-ethylmaleimide, hydrolyzed N-ethylmaleimide and cysteinylation of cysteine. MS1 and MS2 mass accuracies were set to 15 ppm. Peptidoform scoring and match-between-runs were enabled. Results were exported at 5% false discovery rate and filtered to 1% false discovery rate for downstream analysis. A second search with an extended library is described in the Supplementary Methods.

Cysteinylated sites analysis was performed separately. Precursors were filtered at run-specific precursor, protein-group and peptidoform q-values ≤ 0.01 and a library peptidoform q-value ≤ 0.01. Cysteinylated precursors additionally required a post-translational modification site confidence ≥ 0.9. Site occupancy was calculated per sample as the summed DIA-NN quantity of the cysteinylated precursors divided by the summed quantity of all precursors covering the same cysteine (cysteinylated, N-ethylmaleimide, hydrolyzed N-ethylmaleimide and carbamidomethylated forms), without normalization. A site was retained if the cysteinylated form was detected in at least three of five replicates of any condition, the non-cysteinylated forms were detected in at least three samples, the median occupancy across all samples was ≤ 10%, and the retention time was consistent with cysteinylation, i.e. elution before the N-ethylmaleimide and hydrolyzed N-ethylmaleimide forms, at most 1 min after the carbamidomethylated form and no co-elution with another form. Differences in occupancy between conditions were tested with limma on untransformed occupancy values (in percent) as described above, using the same contrasts and Benjamini–Hochberg adjustment, for sites with at least three values per condition.

### Extended spectral library generation and search with fine-tuned predictors

To include all cysteine forms of the cysteine-containing peptides of the ddaPASEF spectral library, the deep-learning predictors of DIA-NN 2.7.0 were fine-tuned on this library. For the training set, 27 of the 62 cysteinylated peptidoforms were removed from the library, either because no other form of the same cysteine was present in the library or because their library retention time was inconsistent with cysteinylation. The retention time, ion mobility and fragmentation predictors were then fine-tuned on this training set (--tune-lib, --tune-rt, --tune-im, --tune-fr). The tuned models were validated on five cysteinylated peptides withheld from training, which gave a median absolute retention time error of 0.9 min, and all five were correctly predicted to elute before the corresponding carbamidomethylated form. With the tuned models, all cysteine forms (with up to one further variable modification) of the 11,607 cysteine-containing peptides of the ddaPASEF library were predicted. Predicted precursors were added to the library if their peptidoform was not already present, every cysteine carried one of the four cysteine modifications (carbamidomethyl, N-ethylmaleimide, hydrolysed N-ethylmaleimide or cysteinyl), and their charge state was present in the ddaPASEF library. This added 54,866 precursors. Predicted retention times and ion mobilities were calibrated to the ddaPASEF library by linear regression (R^2^ = 0.98 and 0.97, respectively). The resulting extended library contained 192,965 precursors.

The 20 diaPASEF runs were searched with the extended library in DIA-NN 2.7.0 using the same settings as the search with the ddaPASEF spectral library. Cysteinylation sites were filtered, quantified, selected and tested exactly as described there. Sites from both searches were matched by gene and cysteine position.

### Cell death assay

To assess cell death, cells were seeded in 12-well plates one day before treatment. After being cultured with 100 µM BSO and other reagents for 72 hours, the cells were trypsinized and collected in 1.7-ml microtubes, washed once with PBS and resuspended in 1 µg ml^−1^ propidium iodide (PI; Sigma P4864-10) in cold PBS. The dead (PI-positive) cells were analysed by FACS using an Attune NxT (Thermo Fisher Scientific) flow cytometer and the FlowJo v10 software.

### Assessment of disulfide stress in dissected tumors

For assessing disulfidptosis-related biomarkers, dissected tumors were lysed in NuPAGE^TM^ LDS sample buffer (Thermo Fisher Scientific, NP0007) without reducing agent. Each sample is then split into two aliquots: one for non-reducing analysis and one for reducing analysis. For the reducing aliquot, β-mercaptoethanol is added to a final concentration of 2%. Both non-reducing and reducing samples are heated at 70°C for 10 min before SDS-PAGE, and 20–30 μg total protein was loaded. Drebrin (Proteintech, 10260-1-AP) and β-actin (Thermo, MA5-11869) oligomerization in non-reducing samples were compared to the samples supplemented with β-mercaptoethanol.

### Compound screen with a fragment-based electrophilic covalent probe library

To identify pharmacological tools that enhance cystine toxicity under conditions of GSH limitation, a compound library screen was performed using the Electrophilic Covalent Probe Library (960 compounds; Enamine, ECPL-960) at the screening facility of the Rudolf-Virchow-Zentrum (RVZ). H460 parental cells were resuspended in DMEM containing 800 μM L-cystine and 500 nM Lip-1, supplemented with either 20 μM BSO or the corresponding vehicle (PBS). Cells were seeded at 1250 cells per well in 384-well plates using an Agilent FX406 liquid-handling system. After overnight incubation, the plates were divided into two treatment conditions: 1 μM erastin co-treatment and solvent (DMSO) control.

Library compounds were dispensed into the wells using an ECHO 525 liquid handler. Each compound was tested at three concentrations: 0.89 μM (low), 5 μM (middle), and 16.7 μM (high). All wells were normalized to a final DMSO concentration of 1%, excluding the control wells for DMSO viability effects.

After 48 hours of incubation, 30 μL of resazurin solution was added to each well using an Agilent FX406 liquid-handling system. The resazurin solution was prepared by diluting a 0.5% stock solution 150-fold in DMEM containing 200 μM L-cystine. After 3 hours of incubation, fluorescence was measured using a Tecan Spark 10M microplate reader (excitation, 550 nm; emission, 600 nm). All data was analyzed using R and Rstudio.

For primary hit identification, data from the intermediate compound concentration (5 μM) were analyzed. Compounds were initially selected if their viability was within the predefined threshold of less than 5 × SD of the control values. Compounds that reduced cell viability in both the presence and absence of erastin were excluded. The remaining compounds identified under the no-erastin condition were subsequently evaluated under the 1 μM erastin treatment condition.

As a second selection criterion, compounds exhibiting a rescue effect by co-treatment of erastin greater than 1.49 and 1.29 were selected as Hit1 and Hit2 molecules respectively. This screening workflow identified 11 Hit1 and 34 Hit2 molecules, yielding 45 candidate compounds in total.

The 45 candidate compounds were subsequently validated using the same experimental setup for five compound concentrations (0.89, 1.6, 2.5, 3.3, and 5 μM). As selection criteria the viability difference between erastin/compound co-treatment and compound treatment alone was used. Six compounds were visually selected showing a concentration dependent delta viability.

### *In vivo TXNDC17*-knockout tumor xenograft model

The xenograft experiments were performed in accordance with a protocol approved by the Institutional Animal Care and Use Committee and Institutional Review Board at The University of Texas MD Anderson Cancer Center. The study is compliant with all relevant ethical regulations regarding animal research. Female athymic nude mice (Foxn1nu/Foxn1nu; 4–6 weeks old; 6 mice per group) were used for the cell line xenograft experiments. Mice were housed under specific-pathogen-free conditions with a 12 h light– 12 h dark cycle. The ambient temperature was 21–23 °C, with 45% humidity and the mice had ad libitum access to water and food. H460 *TXNDC17*^KO^ or its control (NT) cells were resuspended in FBS-free DMEM medium and injected into mice subcutaneously. The tumor volume was calculated according to the equation: volume = 0.5 × length × width^2^. When the tumors had grown to around 50–100 mm^3^ in volume, the mice were randomly assigned to different groups and treated with subcutaneous administration of 360 mg kg^−1^ BSO in 100 µl of saline (vehicle) or vehicle alone every two days. No statistical method was used to pre-determine the sample size. The investigators were not blinded to treatment allocation during experiments or the outcome assessment.

### mRNA sequencing and pathway enrichment analysis

A375 TXNDC17-knockout cells carrying an empty vector (mock) or reconstituted with wild-type TXNDC17 were treated with 100 µM BSO for 36 h. Total RNA was extracted using the Monarch Total RNA Miniprep Kit according to the manufacturer’s instructions. RNA quality was checked using a 5200 Fragment Analyzer with the DNF-471-33 - SS Total RNA 15nt kit (Agilent Technologies). The RQN for all samples was ≥ 9,3. DNA libraries suitable for sequencing were prepared from 200 ng of total RNA with oligo-dT capture beads for poly-A-mRNA enrichment using the NEB mRNA stranded Library Preparation Kit (Illumina) according to manufacturer’s instructions (½ volume). After 12 cycles of PCR amplification, the size distribution of the barcoded DNA libraries was estimated ∼340 bp by electrophoresis on 5200 Fragment Analyzer with the DNF-474-33 - HS NGS Fragment 1-6000bp kit (Agilent Technologies).

Sequencing of pooled libraries, spiked with 1% PhiX control library, was performed at ∼25 million reads/sample in single-end mode with 100 nt read length on the NextSeq 2000 platform (Illumina). Demultiplexed FASTQ files were generated with bcl-convert v4.3.6 (Illumina).

To assure high sequence quality, Illumina reads were quality- and adapter-trimmed via Cutadapt ^31^ v2.5 using a cutoff Phred score of 20 in NextSeq mode, and reads without any remaining bases were discarded (parameters: --nextseq-trim=20 -m 1 -a AGATCGGAAGAGCACACGTCTGAACTCCAGTCAC). Processed reads were subsequently mapped to the human genome (NCBI RefSeq assembly GCF_000001405.40_GRCh38.p14) using STAR ^32^ v2.7.2b with default parameters but including transcript annotations from RefSeq annotation version RS_2025_08 for GRCh38.p14. This annotation was also used to generate read counts on exon level summarized for each gene via featureCounts v1.6.4 from the Subread package ^33^. Multi-mapping and multi-overlapping reads were counted strand-specific and reversely stranded with a fractional count for each alignment and overlapping feature (parameters: -s 2 -t exon -M -O --fraction). The count output was utilized to identify differentially expressed genes using DESeq2 ^34^ v1.24.0. Read counts were normalized by DESeq2 and fold-change shrinkage was conducted by setting the parameter betaPrior to TRUE. Differential expression was assumed at adjusted p-value after Benjamini-Hochberg correction (padj) < 0.05 and |log2FoldChange| ≥ 1. clusterProfiler ^35^ v4.8.1 was used to perform functional enrichment analyses based on Kyoto Encyclopedia of Genes and Genomes (KEGG) pathways and Gene Ontology (GO) terms. The enricher function was used to perform over-representation analysis (ORA) based on lists of significant genes and the GSEA function was applied for gene set enrichment analysis considering the DESeq2 log2FoldChange of all analyzed genes.

### Software and data processing

Graphs were generated using GraphPad Prism v.10 (GraphPad Software) if not stated otherwise. One-way ANOVA followed by Tukey’s multiple-comparisons test was performed with GraphPad Prism v.10 to compare the metabolites in different genotypes and treatments. Gene dependency data were obtained from the Cancer Dependency Map Project at the Broad Institute (DepMap, version 26Q3, https://www.depmap.org/portal), accessed on November 6, 2026.

## Acknowledgments

J.P.F.A. acknowledges the support of from Rudolf Virchow Center, University of Würzburg and additional grants from the Deutsche Forschungsgemeinschaft (DFG), FR 3746/5-1, FR 3746/6-1, CRC/TRR205, INST269/886-1, and TRR 387/1 (514894665); the EU-H2020 (ERC-Consolidator, DeciFERR); and the Deutsche Jose Carreras Leukämie Stiftung (DJCLS 01 R/2022). We thank the Core Unit SysMed at the University of Würzburg for technical support, RNA-seq data generation and analysis. Their work is supported partially by the IZKF at the University of Würzburg (project Z-6).

## Author contributions

C.C.F., M.S. and S.S. developed methodology, performed most of the experiments, analyzed experimental and bioinformatics data, and wrote the original draft. T.N.X.S. performed and analyzed experiments. Q.L. and B.G. supported reagents and infrastructure and performed the in vivo experiments. S.B. developed methodology, performed and analyzed the compound screen. S.O., U.B. and T.A. performed and analyzed targeted sulfur metabolome experiments. G.F. contributed to writing, interpretation and discussion. E.S.J.A. supported CTE probes validation. G.P. and C.B. supported reagents and infrastructure and performed and analyzed MS-based proteomics experiments. J.P.F.A. connived and supervised the project. The final version received input from all authors. All authors contributed to data discussion, interpretation, revision and approved the final version of the manuscript.

## Data, code, and materials availability

The mass spectrometry proteomics data have been deposited to the ProteomeXchange Consortium via the PRIDE ^36^ partner repository with the dataset identifier PXD086652.

## Artificial intelligence

Coding of calculation of S-cysteinylated site occupancy and visualization was supported by Claude (Opus 5.5). Generative artificial intelligence was used to assist with language editing. The authors are responsible for the accuracy and integrity of the final manuscript.

---

**Extended Data Fig. 1.**
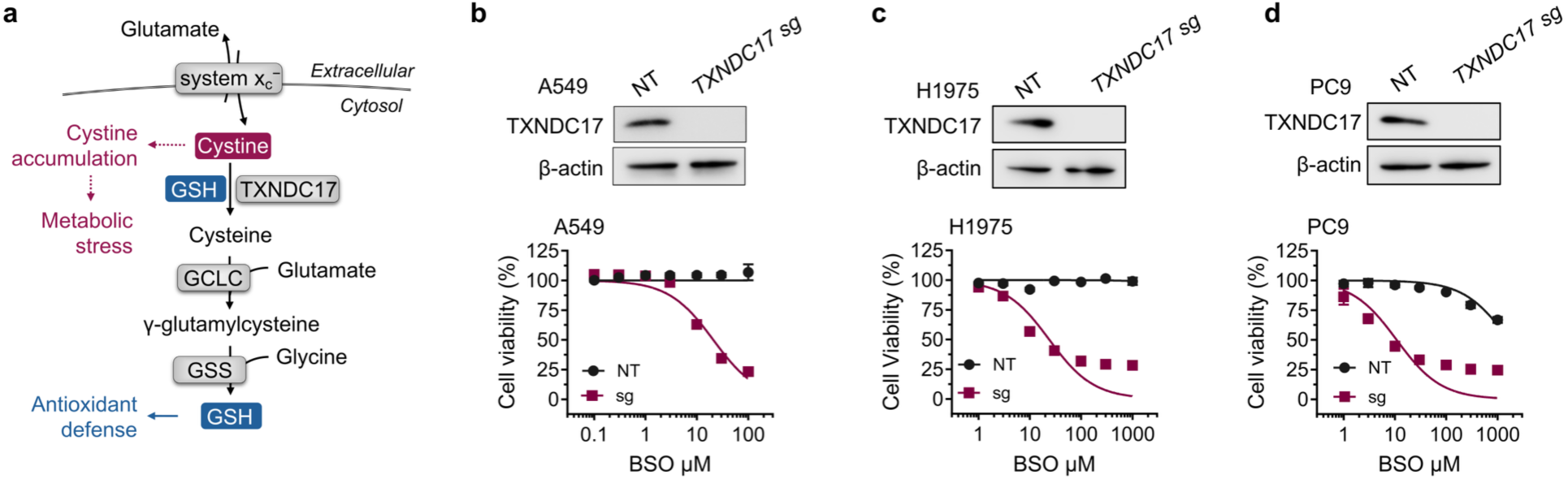
Validation of TXNDC17-dependent survival during GSH depletion. **a.** Cystine uptake and reduction are coupled to glutathione synthesis. System x_c_^−^ imports extracellular cystine in exchange for intracellular glutamate. TXNDC17 and reduced glutathione (GSH) support cytosolic cystine reduction to cysteine. Thioredoxin reductase 1 (TXNRD1) regenerates reduced TXNDC17, and glutathione reductase (GR) reduces glutathione disulfide (GSSG) to GSH. Both reactions use NADPH. The catalytic subunit of glutamate–cysteine ligase (GCLC) joins cysteine and glutamate to form γ-glutamylcysteine, to which glutathione synthetase (GSS) adds glycine to produce GSH. Erastin inhibits system x_c_^−^, whereas buthionine sulfoximine (BSO) blocks the first step of GSH synthesis by inhibiting GCLC. **b-d.** Top: Immunoblots of TXNDC17 and Bottom: Cell viability following BSO treatment in *TXNDC17*-knockout cells and the corresponding non-targeting control cells (NT) in A549 (**b**), H1975 (**c**) and PC9 cells (**d**).

**Extended Data Fig. 2.**
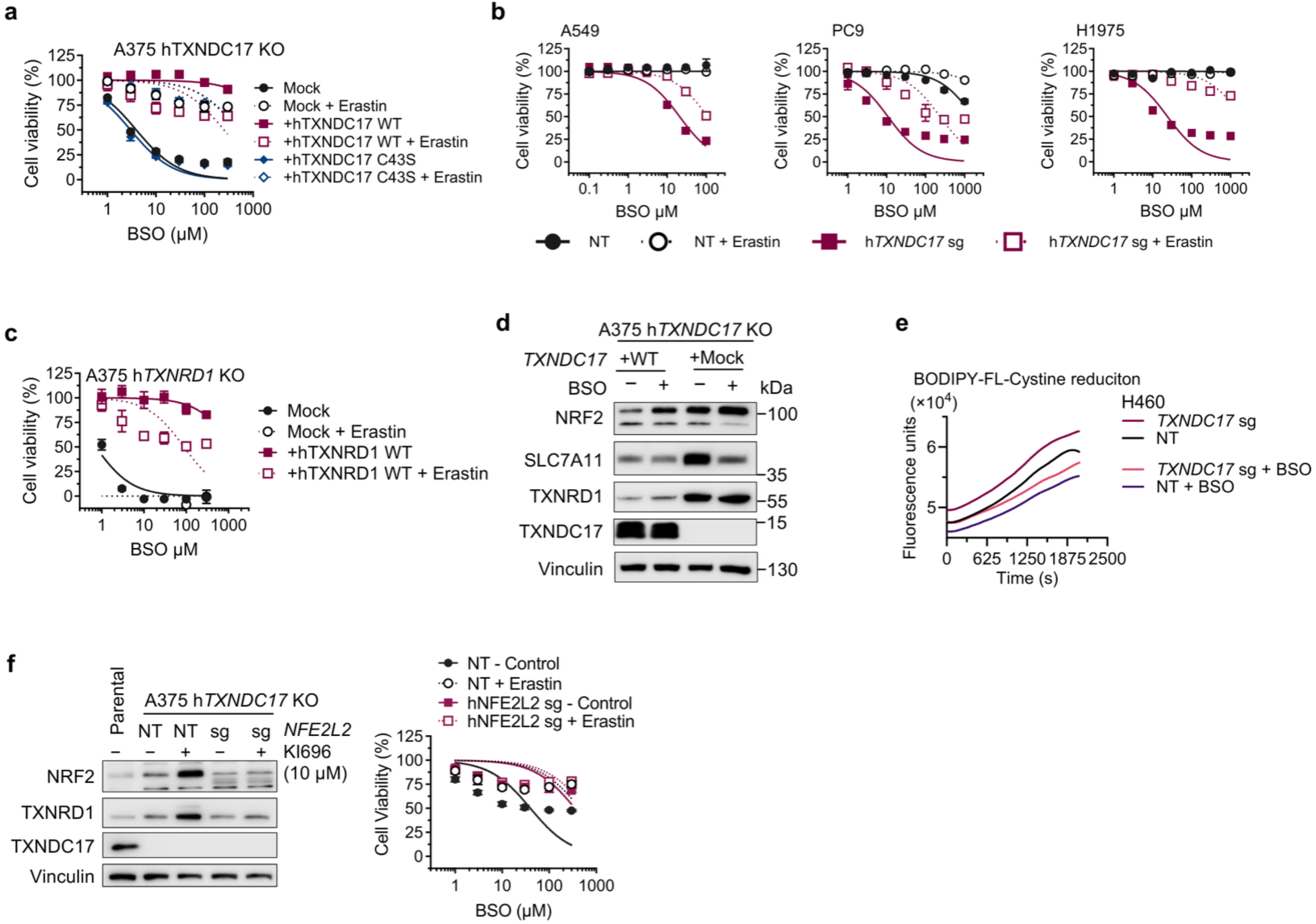
SLC7A11 regulates cytotoxicity in GSH-depleted, TXNDC17-deficient cells. **a.** Cell viability following BSO treatment with 1 µM erastin or vehicle (DMSO) in A375 cells of the indicated genotype. Immunoblots of A375 *TXNDC17*-knockout cells expressing *TXNDC17*-wildtype or mock control cells. Cells were treated with 50 µM BSO for 36 hours. **b.** Cell viability following BSO treatment with 1 µM erastin or vehicle (DMSO) in A549 (left), PC9 (middle), and H1975 (right). **c.** Cell viability performed as described in Extended Data Fig. 2a using A375 *TXNRD1*-knockout cells expressing *TXNRD1*-wildtype or mock control cells. **d.** Immunoblots of A375 *TXNDC17*-knockout cells expressing *TXNDC17*-wildtype or mock control cells. Cells were treated with 50 µM BSO for 36 hours. **e.** Kinetic analysis of cystine reduction by *TXNDC17*, monitored using BODIPY™-FL L-cystine. H460 *TXNDC17*-knockout cells and NT control cells were incubated with 10 µM BSO or vehicle (PBS) for 24 h before the assay. **f.** Left: Immunoblots of A375 cells with single or double knockout of *TXNDC17* and *NFE2L2*. Cells were treated with 10 µM KI696 for 48h to induce NRF2 activation. Right: Cell viability following BSO treatment in the presence of 1 µM erastin or vehicle (DMSO).

**Extended Data Fig. 3.**
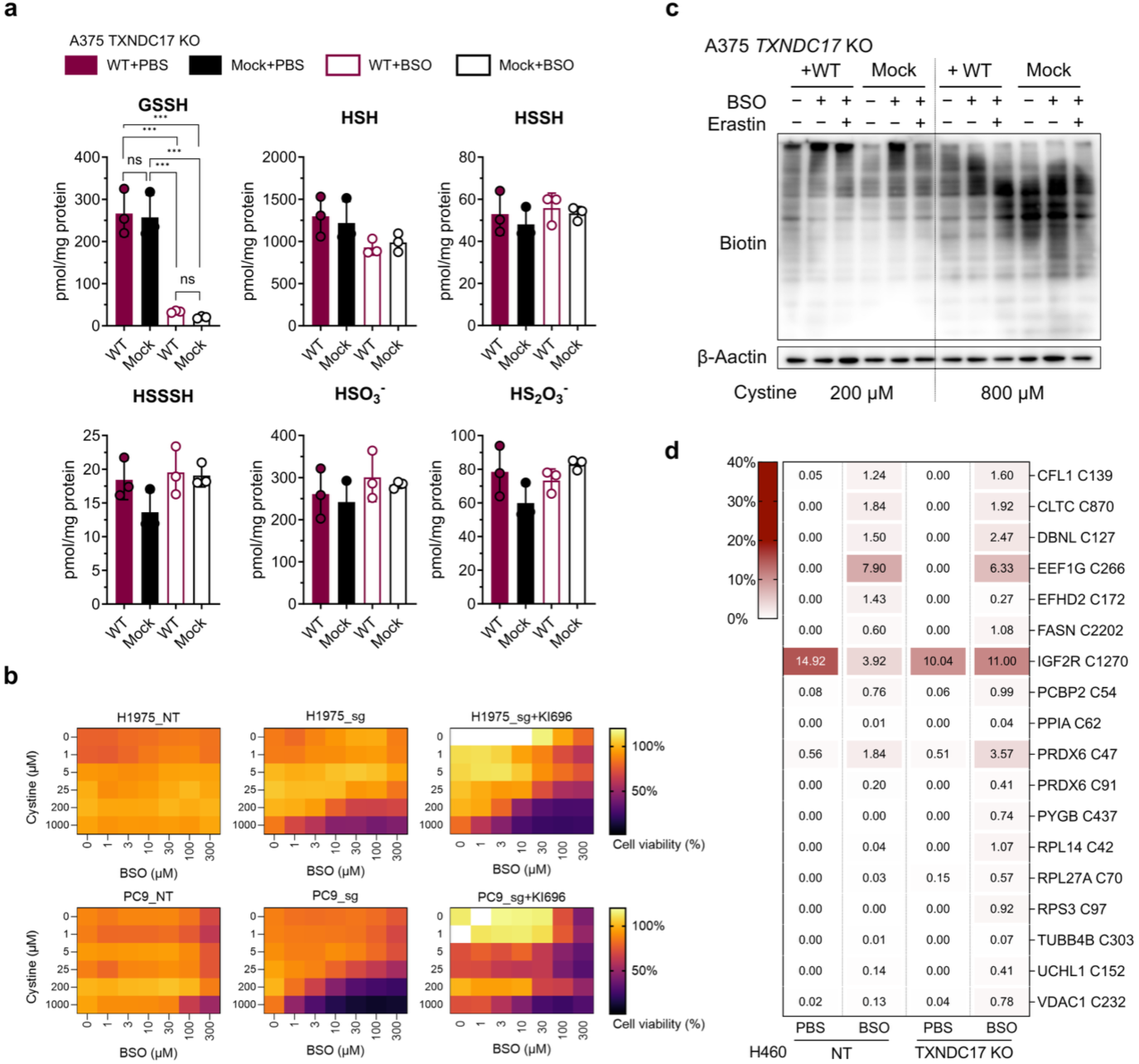
Cystine-dependent cytotoxicity and protein cysteinylation in GSH-depleted, TXNDC17-deficient cells. **a.** Quantification of sulfur metabolites by LC–MS/MS. A375 *TXNDC17*-knockout cells expressing wild-type TXNDC17 or mock control cells were incubated with 50 µM BSO or vehicle (PBS) for 24 h. GSSH, glutathione persulfide; HSH, hydrogen sulfide; HSSSH, hydrogen trisulfide; HSO_3_^−^, Sulfite; HS_2_O_3_^−^, thiosulfate. Data are presented as mean ± SD. Statistical analysis was performed using one-way ANOVA followed by Tukey’s multiple-comparisons test. ns, not significant (P > 0.05); ***P ≤ 0.001. **b.** Dose-dependent cystine toxicity under conditions of glutathione depletion. Cell viability was assessed in *TXNDC17-*knockout (sg) and the corresponding non-targeting control (NT) cells from the indicated human non-small lung cell cancer cell lines. The cells were treated with BSO in medium containing different concentrations of L-cystine. KI696 10 µM pre-treatment was performed 72h before BSO exposition. **c.** Analysis of protein cysteinylation using N-biotinyl-L-cysteine in A375 *TXNDC17*-knockout cells reconstituted TXNDC17-wildtype or mock control cells. Cells were incubated for 24 h in medium containing 200 µM or 800 µM cystine and treated with PBS or 100 µM BSO in combination with 1 µM erastin or vehicle (DMSO), as indicated. **d.** Expanded-library reanalysis of the DIA dataset shown in Fig. 2j, displaying additional S-cysteinylated site occupancies across NT and *TXNDC17*-knockout cells with 50 µM BSO or PBS for 24 h. All cells were additionally treated with 500 nM Lip-1.

**Extended Data Fig. 4.**
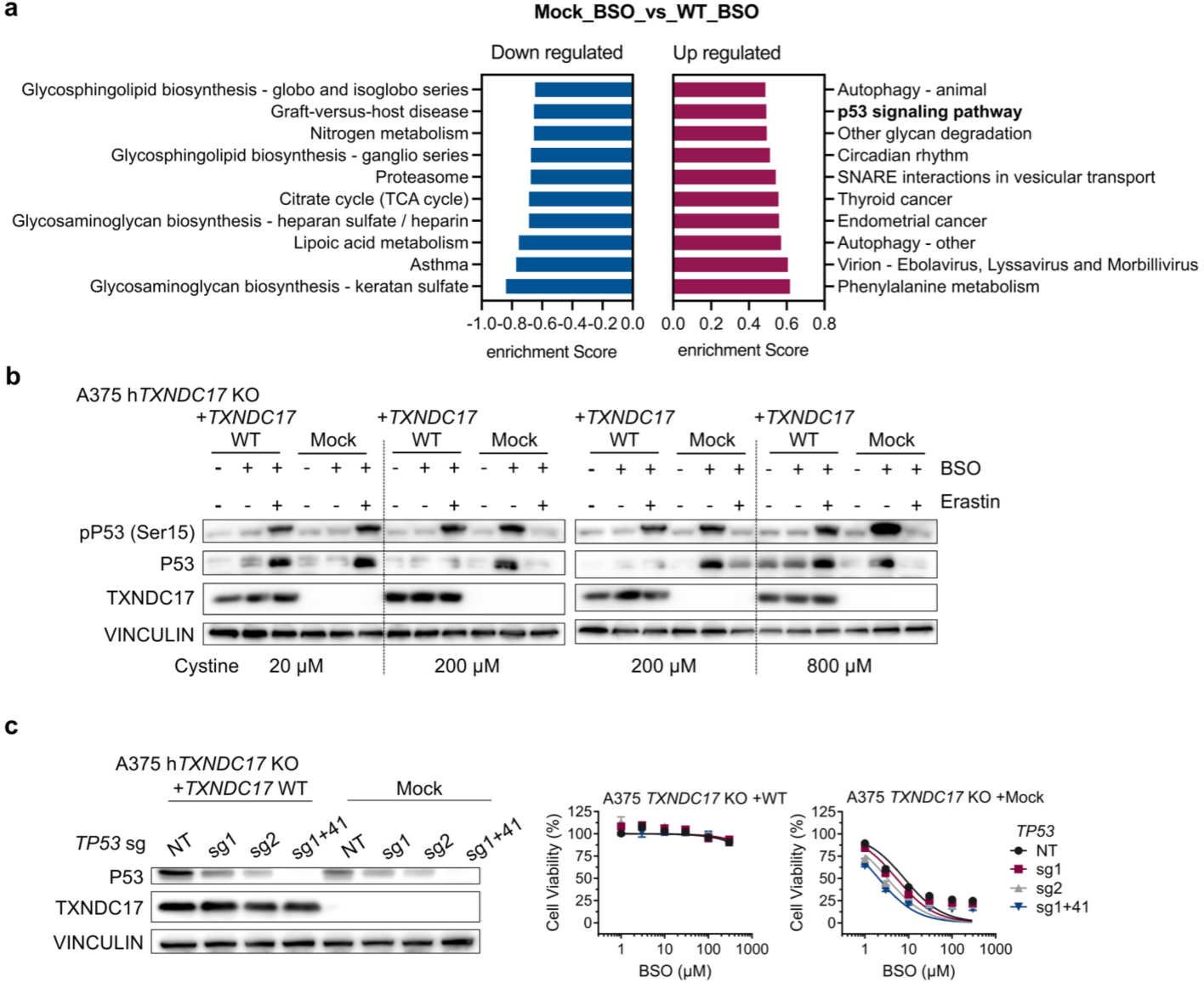
Exploratory profiling of stress responses to combined TXNDC17 loss and GSH depletion. **a.** Pathway enrichment analysis of the transcriptional response to 100 µM BSO-treated A375 *TXNDC17*-knockout cells carrying mock or WT TXNDC17 addback for 36 h. RNA-seq profiles were compared between BSO-treated *TXNDC17*-knockout cells with mock addback and BSO-treated knockout cells reconstituted with wild-type TXNDC17. Negative (blue) and positive (magenta) enrichment scores indicate pathways associated with lower and higher transcript abundance, respectively, in mock-addback cells relative to wild-type TXNDC17–reconstituted cells. The p53 signaling pathway was among the positively enriched pathways. **b.** Immunoblots of p53 phosphorylated at Ser15, total p53, TXNDC17, and vinculin in A375 *TXNDC17*-knockout cells carrying WT TXNDC17 or mock addback, cultured with the indicated extracellular L-cystine concentrations and treated with 50 µM BSO and/or 1 µM erastin as shown for 36 h. The figure includes two blocks labeled 200 µM cystine as membrane exposition control. **c.** Left: Immunoblots of P53 and TXNDC17 in A375 *TXNDC17*-knockout cells reconstituted TXNDC17-wildtype or mock control cells plus different sgRNA (sg1, sg2 and sg1+41 combination) targeting *TP53*. Right: Cell viability following BSO treatment in the same cells.

**Extended Data Fig. 5.**
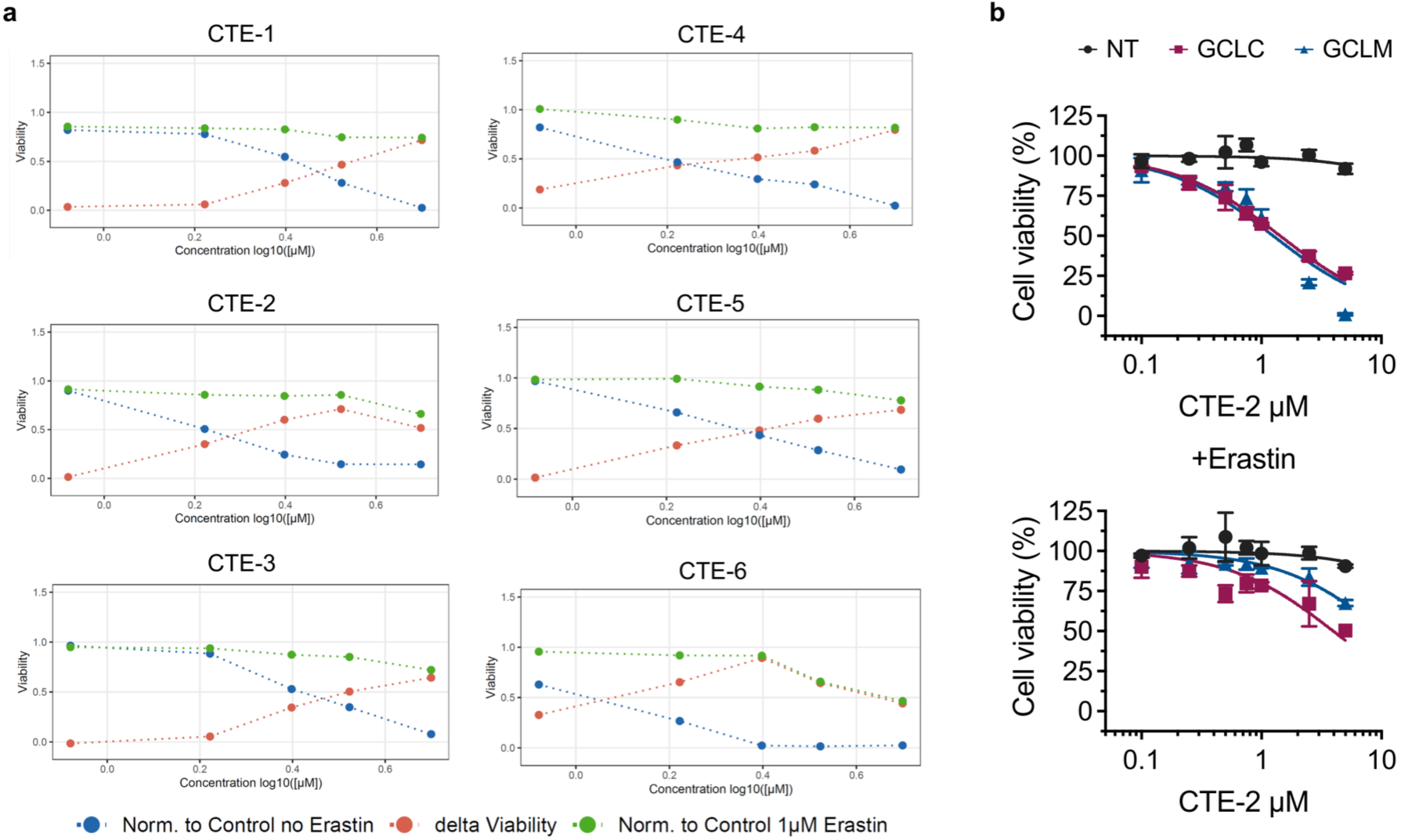
Validation of cystine-toxicity enhancers (CTE) **a.** Selected candidates from the primary screen were retested under the same screening conditions at five concentrations (0.89, 1.6, 2.5, 3.3, and 5 μM). The X-axis represents fragments concentrations, and the Y-axis represents cell viability normalized to that of DMSO-treated control wells. The blue and green lines indicate cell viability in the absence or presence of 1 μM erastin, respectively. The red line indicates the difference in cell viability between the two conditions (delta viability). The six top-ranked probes are shown and designated as cystine-toxicity enhancers (CTE-1 to CTE-6). **b.** Cell viability following treatment with CTE-2 in H460 GCLC-knockout and GCLM-knockout cells in the presence of DMSO (top) or 1 µM erastin (bottom) for 48 h.

**Extended Data Table 1.**
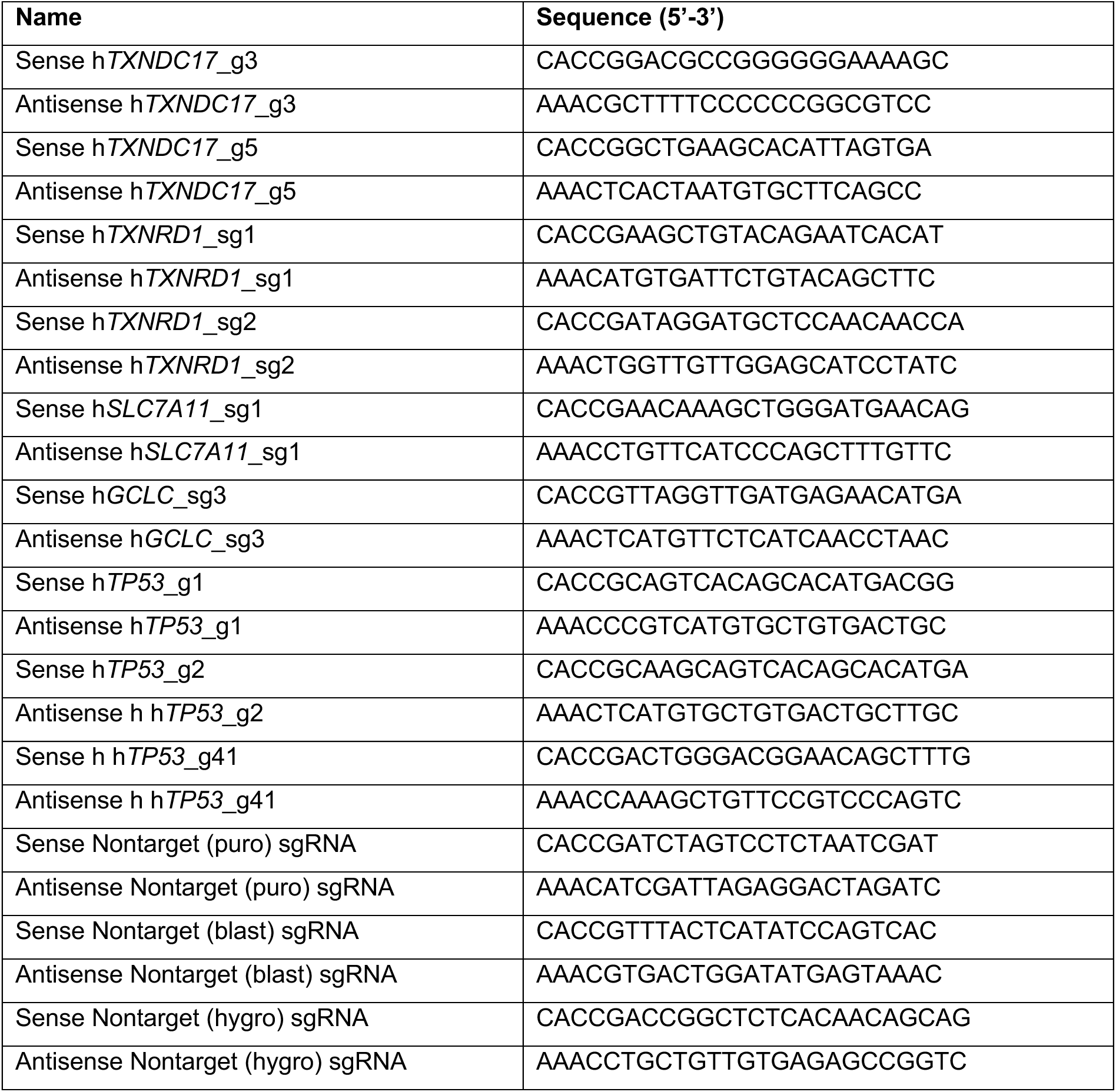
Oligonucleotide sequences used for CRISPR Cas9-mediated gene knockout.

**Extended Data Table 2.**
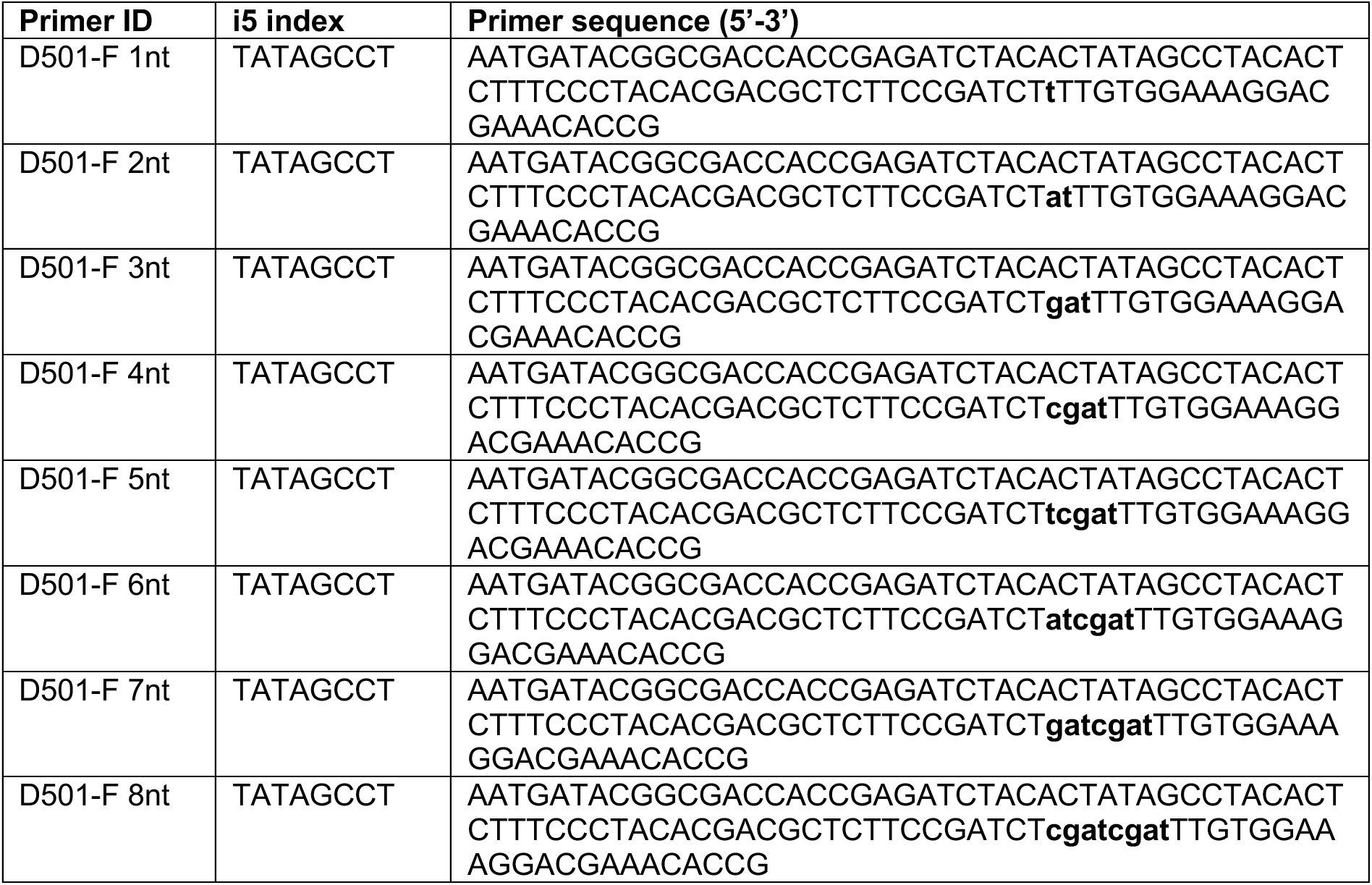
Sequences of staggered forward primers used for Illumina sequencing library preparation.

**Extended Data Table 3.**
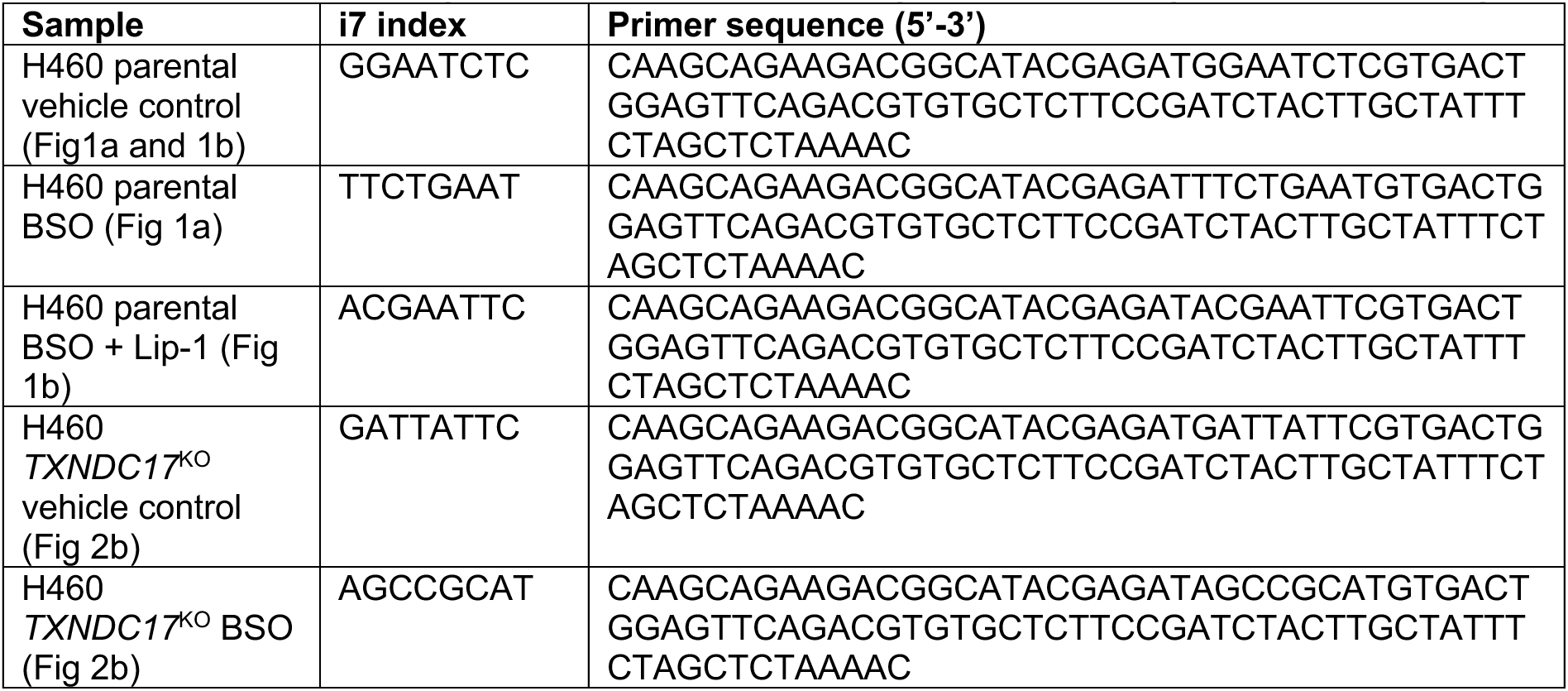
Sequences of indexed reverse primers for multiplexed Illumina sequencing.

**Extended Data Table 4.**
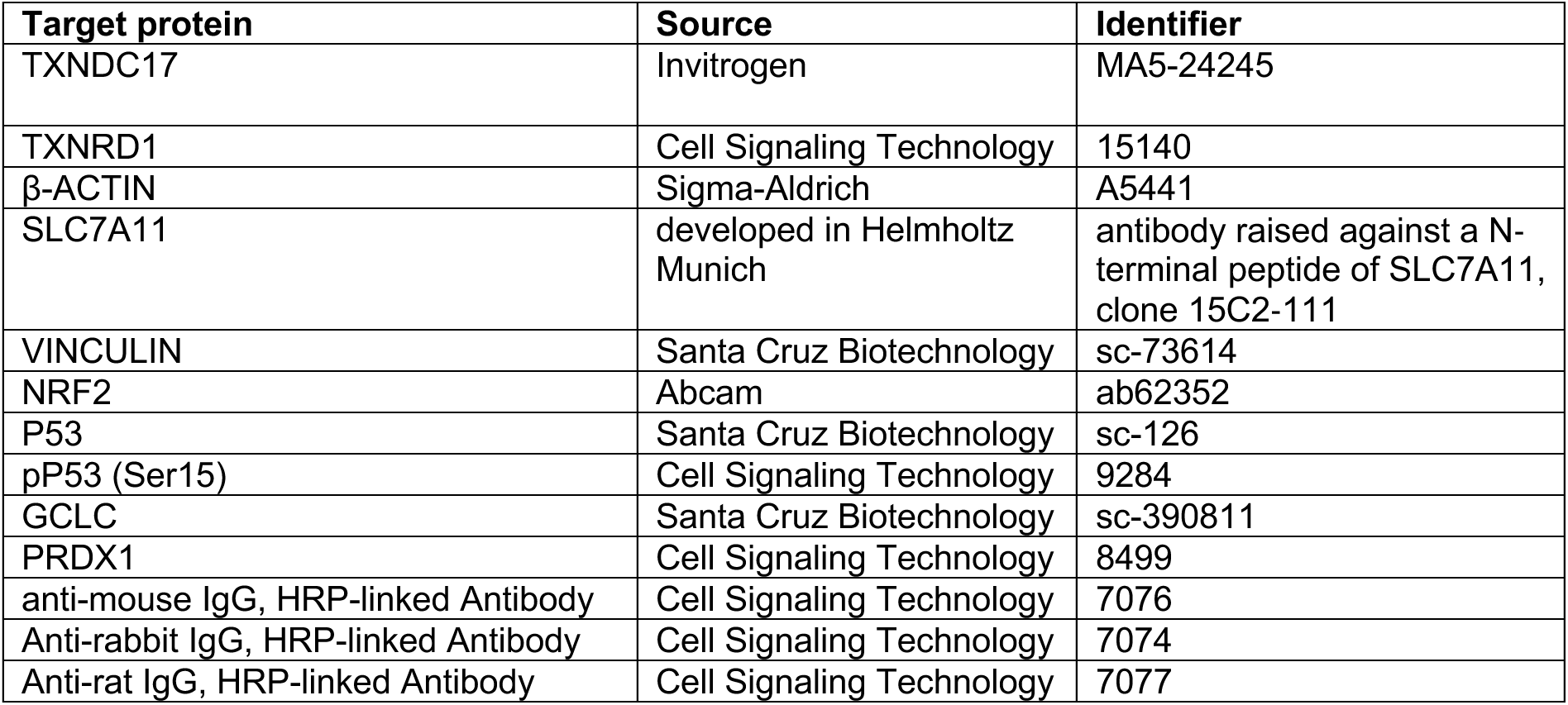
Antibody used in this study.

**Extended Data Table 5.** Data independent acquisition-PASEF isolation window scheme.

| **Ramp** | **Window 1: m/z (1/K0)** | **Window 2: m/z (1/K0)** | **Window 3: m/z (1/K0)** |
|---|---|---|---|
| 1 | 400–425 (0.64–0.83) | 600–625 (0.83–1.01) | 800–825 (1.01–1.37) |
| 2 | 425–450 (0.64–0.85) | 625–650 (0.85–1.04) | 825–850 (1.04–1.37) |
| 3 | 450–475 (0.64–0.87) | 650–675 (0.87–1.06) | 850–875 (1.06–1.37) |
| 4 | 475–500 (0.64–0.90) | 675–700 (0.90–1.09) | 875–900 (1.09–1.37) |
| 5 | 500–525 (0.64–0.92) | 700–725 (0.92–1.11) | 900–925 (1.11–1.37) |
| 6 | 525–550 (0.64–0.94) | 725–750 (0.94–1.13) | 925–950 (1.13–1.37) |
| 7 | 550–575 (0.64–0.97) | 750–775 (0.97–1.16) | 950–975 (1.16–1.37) |
| 8 | 575–600 (0.64–0.99) | 775–800 (0.99–1.18) | 975–1,000 (1.18–1.37) |

## REFERENCES

1. Berndt, C., Lillig, C.H. & Flohe, L. Redox regulation by glutathione needs enzymes. Front Pharmacol 5, 168 (2014).

2. Dixon, S.J. & Olzmann, J.A. The cell biology of ferroptosis. Nat Rev Mol Cell Biol 25, 424–442 (2024).

3. Dixon, S.J. et al. Ferroptosis: an iron-dependent form of nonapoptotic cell death. Cell 149, 1060–72 (2012).

4. Yang, W.S. et al. Regulation of ferroptotic cancer cell death by GPX4. Cell 156, 317–331 (2014).

5. Friedmann Angeli, J.P., et al. Inactivation of the ferroptosis regulator Gpx4 triggers acute renal failure in mice. Nat Cell Biol 16, 1180–91 (2014).

6. Lushchak, V.I. Glutathione homeostasis and functions: potential targets for medical interventions. J Amino Acids 2012, 736837 (2012).

7. Xiong, Y., Uys, J.D., Tew, K.D. & Townsend, D.M. S-glutathionylation: from molecular mechanisms to health outcomes. Antioxid Redox Signal 15, 233–70 (2011).

8. Ward, N.P. et al. Mitochondrial respiratory function is preserved under cysteine starvation via glutathione catabolism in NSCLC. Nat Commun 15, 4244 (2024).

9. Hecht, F. et al. Catabolism of extracellular glutathione supplies cysteine to support tumours. Nature 653, 933–941 (2026).

10. Sato, H., Tamba, M., Ishii, T. & Bannai, S. Cloning and expression of a plasma membrane cystine/glutamate exchange transporter composed of two distinct proteins. J Biol Chem 274, 11455–8 (1999).

11. Wu, G. et al. Glutathione metabolism and its implications for health. J Nutr 134, 489–92 (2004).

12. Sasaki, H. et al. Electrophile response element-mediated induction of the cystine/glutamate exchange transporter gene expression. J Biol Chem 277, 44765–71 (2002).

13. Zhang, D.D., Lo, S.C., Cross, J.V., Templeton, D.J. & Hannink, M. Keap1 is a redox-regulated substrate adaptor protein for a Cul3-dependent ubiquitin ligase complex. Mol Cell Biol 24, 10941–53 (2004).

14. Sayin, V.I. et al. Activation of the NRF2 antioxidant program generates an imbalance in central carbon metabolism in cancer. Elife 6(2017).

15. Griffith, O.W. & Meister, A. Potent and specific inhibition of glutathione synthesis by buthionine sulfoximine (S-n-butyl homocysteine sulfoximine). J Biol Chem 254, 7558–60 (1979).

16. Liu, X. et al. Actin cytoskeleton vulnerability to disulfide stress mediates disulfidptosis. Nat Cell Biol 25, 404–414 (2023).

17. Doll, S. et al. FSP1 is a glutathione-independent ferroptosis suppressor. Nature 575, 693–698 (2019).

18. Mandal, P.K. et al. System x(c)- and thioredoxin reductase 1 cooperatively rescue glutathione deficiency. J Biol Chem 285, 22244–53 (2010).

19. Pader, I. et al. Thioredoxin-related protein of 14 kDa is an efficient L-cystine reductase and S-denitrosylase. Proc Natl Acad Sci U S A 111, 6964–9 (2014).

20. Yan, X. et al. Inhibition of Thioredoxin/Thioredoxin Reductase Induces Synthetic Lethality in Lung Cancers with Compromised Glutathione Homeostasis. Cancer Research 79, 125–132 (2019).

21. Marti-Andres, P. et al. TRP14 is the rate-limiting enzyme for intracellular cystine reduction and regulates proteome cysteinylation. EMBO J 43, 2789–2812 (2024).

22. Sastre, J. et al. Integrated roles of the thioredoxin-related protein of 14 kDa, TRP14, in sulfur amino acid metabolism, protein posttranslational modifications and signaling. Free Radic Biol Med 257, 545–560 (2026).

23. Jiang, L. et al. Ferroptosis as a p53-mediated activity during tumour suppression. Nature 520, 57–62 (2015).

24. Schmidt, E.E. et al. Cystine C-S bond cleavage fuels cysteine production under disulfide reductase deficiency. Nat Chem Biol (2026).

25. Tang, M. et al. Inhibition of thioredoxin reductase 1 sensitizes glucose-starved glioblastoma cells to disulfidptosis. Cell Death Differ 32, 598–612 (2025).

26. Li, W. et al. MAGeCK enables robust identification of essential genes from genome-scale CRISPR/Cas9 knockout screens. Genome Biol 15, 554 (2014).

27. Akaike, T. et al. Cysteinyl-tRNA synthetase governs cysteine polysulfidation and mitochondrial bioenergetics. Nat Commun 8, 1177 (2017).

28. Matsunaga, T. et al. Supersulphides provide airway protection in viral and chronic lung diseases. Nat Commun 14, 4476 (2023).

29. Hughes, C.S. et al. Single-pot, solid-phase-enhanced sample preparation for proteomics experiments. Nat Protoc 14, 68–85 (2019).

30. Demichev, V., Messner, C.B., Vernardis, S.I., Lilley, K.S. & Ralser, M. DIA-NN: neural networks and interference correction enable deep proteome coverage in high throughput. Nat Methods 17, 41–44 (2020).

31. Martin, M. Cutadapt removes adapter sequences from high-throughput sequencing reads. EMBnet.journal 17(2011).

32. Dobin, A. et al. STAR: ultrafast universal RNA-seq aligner. Bioinformatics 29, 15–21 (2013).

33. Liao, Y., Smyth, G.K. & Shi, W. featureCounts: an efficient general purpose program for assigning sequence reads to genomic features. Bioinformatics 30, 923–30 (2014).

34. Love, M.I., Huber, W. & Anders, S. Moderated estimation of fold change and dispersion for RNA-seq data with DESeq2. Genome Biol 15, 550 (2014).

35. Wu, T. et al. clusterProfiler 4.0: A universal enrichment tool for interpreting omics data. Innovation (Camb) 2, 100141 (2021).

36. Perez-Riverol, Y. et al. The PRIDE database at 20 years: 2025 update. Nucleic Acids Res 53, D543–D553 (2025).

